# Synaptation − a missing concept in the science of evolved complexity

**DOI:** 10.64898/2026.01.01.697299

**Authors:** Joseph P. Bielawski

## Abstract

The concept of synaptation is introduced to explain a distinct evolutionary origin of selected effects, in which natural selection operates through emergent cross-level interactions within collectives lacking shared inheritance. The traditional framework excludes such origins from the adaptation–exaptation taxonomy. Yet individual organism fitness is often mediated by collective, non-linear interactions within systems that cannot themselves sustain heredity. To address this, multilevel selection (MLS) theory is extended in the form of synaptive MLS, a process through which heterogeneous task groups evolve emergent traits, called synaptations, whose evolutionary origin lies in irreducible, cross-level selective covariance. Synaptive MLS differs from trait-group selection because collective function depends on causal entanglement among heterogenous entities with diverse fates, rather than a common trait that links entities to a shared fate. This framework applies to conspecific divisions of labor, multi-species consortia and holobiont integration, where coordination of distinct roles yields emergent fitness effects. Synaptation recognizes how natural selection can generate novel emergent functions by coordinating adaptive, exaptive, and neutral organism-level traits whose lower-level roles scaffold higher-level selected effects. Synaptation and adaptation define the endpoints of a continuum of selected-effect complexity. Recognition of this continuum extends the Darwinian paradigm to adaptive complexity in systems that fail classical heredity requirements and offers an evolutionary lens for understanding relational phenomena such as eubiosis, dysbiosis, and rebiosis in holobionts.

**Significance Statement:** Natural selection is typically understood to require heredity, yet some complex and functionally integrated biological systems in nature lack a mechanism for collective reproduction. This project introduces *synaptation*, a theoretical framework that recognizes a distinct evolutionary origin of selected effects, in which fitness consequences arise through non-additive, cross-level interactions rather than through collective reproduction. Synaptive multilevel selection explains how functionally integrated traits can originate by natural selection in non-reproducing collectives, bridging molecular, ecological, and social forms of cooperation. By revealing how coordination of lower-level traits can scaffold the emergence of higher-level selected effects, this framework extends the reach of evolutionary theory and offers new insight into the eco-evolutionary dynamics that sustain collective health, stability, and resilience in multi-species systems.

## 1. Introduction

Darwin’s theory of natural selection is the foundation for understanding how biological complexity evolved to serve a specific function. Darwin’s theory is concisely captured by “Lewontin’s recipe,” which defines natural selection as requiring a population of individuals exhibiting (i) variation, (ii) fitness differences, and (iii) heredity (Lewontin 1970). According to this view, a trait has a selected-effect function only if it evolved within a lineage that satisfies all three of Lewontin’s criteria and it also currently serves the function that was originally selected (Gould and Vrba 1982; Brunet et al. 2021). Complex adaptations evolve over successive generations through integration by natural selection of multiple interdependent components. If all the components were fine-tuned by natural selection for their contribution to the original selected-effect function, then the complex trait is easily classified as an adaptation. However, the status of such a trait as an adaptation can be problematic when its fitness-effecting role changes over time (Gould and Vrba 1982), when it is assembled from several components that individually serve distinct roles (Brunet et al. 2021), or when it manifests at a level that does not meet Lewontin’s heredity criterion (Lean et al. 2022; Doolittle 2024a).

In their seminal paper, Gould & Vrba (Gould and Vrba 1982) described how a flawed taxonomy of fitness-related traits constrained evolutionary reasoning and fostered mistaken ideas about what constitutes a selected-effect function. To correct this, Gould & Vrba (Gould and Vrba 1982) expanded the taxonomy of evolved trait categories by proposing the term *exaptation* for all those traits that currently affect fitness but were *not* built by selection for their current role (Table 1). Exaptation introduced “evolutionary cooption” as an important process. Evolutionary cooption recognizes that biological complexity often arises by repurposing pre-existing genes, genetic systems, or traits for *new roles*. Thus, any trait that originally evolved through selection for a particular function, or having evolved no prior function, becomes an *exaptation* once it is coopted for a new purpose. To better analyze the origin of complex traits composed of parts with diverse evolutionary histories, Brunet et al. (Brunet et al. 2021) formally separated two forms of exaptation described by Gould and Vrba (Gould and Vrba 1982). Here I follow Gould and Vrba (Gould and Vrba 1982) and Brunet et al. (Brunet et al. 2021) by adopting an expanded taxonomy of adaptation/exaptation that distinguishes exaptation^A^, the cooption of previously adaptive traits, from exaptation^N^, the cooption of previously neutral traits. This taxonomy (Table 1) is essential to understanding how a complex trait can emerge from heterogeneous components, each with different selective origins at a single level (Brunet et al. 2021). This classification system remains largely unexamined for traits that function in conspecific or multi-species collectives (Lloyd and Gould 1999). One central objective of this paper is to cover this blind spot.

**Table 1.** A taxonomy of fitness-related traits.

| Character | Status | Co-option | Semantic Category |
| --- | --- | --- | --- |
| Aptation | A trait that currently enhances fitness, making no claims about its evolutionary origin. A super-category that includes adaptation and exaptation. | unspecified | unspecified |
| Nonaptation | A trait that currently does not affect fitness, making no claims about its evolutionary origin. | not at present | neutral |
| Adaptation | A trait that currently enhances fitness *and* was built by natural selection for the use it currently serves. | no | function |
| Exaptation<sup>A</sup><br>(from adaptation) | A trait that was built by natural selection for a particular function (i.e., adaptive at the time of its origin) *and was coopted* for a new use that currently enhances fitness. | yes | effect |
| Exaptation<sup>N</sup><br>(from nonaptation) | A trait that was previously nonaptive *and was coopted* for a new use that currently enhances fitness. | yes | effect |

Multi-species microbial communities can exhibit complex relational traits, analogous to those seen in multicellular organisms (Ereshefsky and Pedroso 2015; Bosch and Miller 2016). For instance, multi-species biofilms exhibit levels of organization, physiological integration and cooperation that resemble those of other autonomous biological entities (Costerton et al. 1995; Marsh and Bowden 2000; O’Malley 2014), prompting some to interpret those features as adaptations at the collective level (De la Fuente-Núñez et al. 2013; Ereshefsky and Pedroso 2015). Similar claims have been made about holobionts—eukaryotic hosts and their associated microbial communities—suggesting that they function as units of selection capable of evolving wholistic adaptations at that level (Zilber-Rosenberg and Rosenberg 2008; Rosenberg and Zilber-Rosenberg 2018). Recent research in community genetics employed clever experimental approaches to support the notion that entire eukaryotic communities could evolve relational adaptations if there existed sufficient levels of community heritability (Whitham et al. 2020). However, interpreting multi-species complexity as adaptive remains contentious. Such collective traits often lack the level of faithful inheritance required by Lewontin’s recipe, casting doubt on their status as canonical units of selection (Lean et al. 2022; Doolittle 2014; Doolittle and Inkpen 2018; Doolittle 2024a).

Collective-level selection is possible, however, even when the collectives are transient, defined by their interactions, and do not have unified inheritance (Wilson 1980; Griesemer 2014). Multi-level selection (MLS) theory shows that selection can change the among-group distribution of a collective trait when it affects the reproductive fitness of lower level individuals (Damuth and Heisler 1988; Okasha 2006). A mechanism for collective trait transmission is not required as long as there is inheritance at some lower level (Damuth and Heisler 1988; Okasha 2006), even for multi-species collectives (Lean et al. 2022; Lean and Jones 2023; Doolittle 2024a). A distinct line of theorizing goes even further and argues that persistence can support evolutionary success in the absence of reproduction, a view developed formally by Bouchard ((Bouchard 2014)) and Bourrat ((Bourrat 2014; 2015)) and later applied by Doolittle and colleagues to clades, symbiotic communities, and planetary-scale biospheres (Doolittle 2014; Doolittle and Booth 2017; Doolittle and Inkpen 2018; Papale and Doolittle 2024). These cases expose a boundary in the taxonomy (Table 1) concerning the evolutionary origin of selected effects: according to Lewontin’s recipe, adaptations cannot belong to what fails to reproduce with heredity (Williams 1966; Maynard Smith 2002; Godfrey-Smith 2009).

As will be shown later, the current taxonomy of fitness-related traits does not recognize the origin of selected effects from emergent cross-level interactions in systems lacking collective heredity. This is despite increasing evidence for lower-level genetic contributions to emergent collective fitness. When genes mediate how one individual alters the phenotype and fitness of another member of the same species (e.g., Wolf et al. 1998; Santostefano et al. 2017; Jaffe et al. 2020), they are said to have indirect genetic effects (IGEs). When genes mediate fitness consequences exchanged across species boundaries (Whitham et al. 2020; Genung et al. 2013; Rebar and Rodríguez 2014; Kelly 2018), they are said to have interspecies indirect genetic effects (IIGEs). IGEs and IIGEs facilitate relational fitness, yet the associated non-additive traits cannot qualify as collective-level adaptations. These observations, together with Griesemer’s (Griesemer 2014) account of “task groups” as heterogeneous collectives that perform coordinated functions without shared heredity, reveal a conceptual gap in how multi-level fitness effects are described. Failure to name distinct origins of fitness effects can lead to their systematic neglect in evolutionary explanations, as seen in earlier conceptual gaps that once obscured phenomena like inclusive fitness (Hamilton 1964) and exaptation (Gould and Vrba 1982).

Queller (Queller 2020) argued that many selective processes involve causal dependencies that span organizational levels, a challenge he termed the ‘Gouldian knot’. Classical MLS frameworks, including contextual analysis and the standard MLS1 partition, assume that selection can be decomposed additively into independent individual- and group-level components. This assumption reinforces the systematic blind spot: when collective function is built from non-additive, configuration-dependent interactions, additive decompositions obscure the cross-level causal structure of fitness. IGEs, IIGEs, and task group effects exemplify this problem, generating fitness consequences that are neither strictly individual nor fully collective. In such systems the trait–fitness covariance can remain entangled across levels even after all additive group descriptors (*e.g*., group means or other linear aggregates) are included. Because evolutionary theory has relied so heavily on additive partitions, our existing taxonomies have never required a category for traits whose fitness consequences are irreducibly cross-level and causally entangled.

To address that gap, this paper introduces the term *synaptation*, which denotes traits whose selective covariance is inherently cross-level and cannot be reduced to individual- or group-level variation alone. The argument for synaptation in framed four steps: first, the general form of natural selection is reviewed; second, the relational blind spots of classical MLS1 are clarified; third, synaptive MLS is developed using a causal-graph framework; and fourth, the biological and conceptual implications are explored. Synaptation is situated as one endpoint in a continuum of selected-effect complexity, with adaptation defining the opposite end. By combining the Price formulation for natural selection with causal modelling, *synaptive MLS* is developed as an extension of selection theory that describes how heterogeneous contributors within *task groups* generate fitness that is irreducible to any single level. Synaptive MLS represents a distinct form of multi-level causation, which applies to both single-species and multi-species MLS. It differs from classical *trait-group* selection (Wilson 1980) because synaptive function arises from non-linear coordination among entities with distinct roles and evolutionary fates. Synaptive MLS explains how interactions within task groups, facilitated by IGEs or IIGEs, can evolve under natural selection even in the absence of group-level heredity. The conceptual framework is illustrated through biological cases involving divisions of labor and simple numerical examples provided in appendices. The concept enhances our understanding of eubiosis, dysbiosis, and rebiosis as evolutionary expressions of functional integration across levels and lineages.

## 2. A general description of evolution by natural selection

The Price equation (Price 1970) is considered by many to be the most general account of evolution by natural selection (*e.g*., (Rice 2004; Gardner 2020)), and it is reviewed here as a foundation for multi-level selection theory. The Price equation is a *mathematical identity* derived from the definitions of expectation and covariance; as such, it does not by itself specify causal mechanisms, directions of causation, or levels of selection.

Consider a population of parents, each with a phenotypic value *z_i_*. A parent’s number of offspring is its fitness (*ɷ_j_*). Under only this bookkeeping, the Price equation describes evolutionary change as:

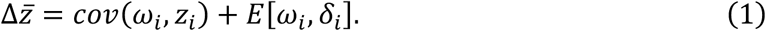

The first term on the right is the selective covariance, highlighting that natural selection cannot occur without covariance between fitness and phenotype. The second term represents an explicit separation of all non-selective processes that can also change the phenotype (*e.g*., mutation and other sources of transmission bias). Importantly, non-zero covariance alone does not establish a causal role for the phenotype (Rice 2004; Okasha 2006); interpretation of the selective covariance requires additional assumptions about how fitness is generated.

Under standard assumptions, the Price equation recovers classical results such as the breeder’s equation, revealing that organism-level natural selection acts on additive, heritable effects on phenotype (SI Appendix 1). This special case illustrates how the general Price framework reduces to familiar models when fitness depends linearly on individual traits and those effects are reliably transmitted across generations. However, this reduction depends on strong assumptions about additivity and causal independence. When fitness depends on interaction-mediated or configuration-dependent effects, these assumptions may fail, and the interpretation of covariance terms becomes ambiguous. Thus, while the Price equation works as a general accounting of evolutionary change, it does not determine whether a given covariance reflects a genuine causal process at a particular level of biological organization. Identifying the causal structure underlying trait–fitness covariance requires additional criteria, particularly criteria that determine when cross-level effects are reducible to additive summaries, which become important in interaction-structured systems.

## 3. MLS1 and the problem of causal decomposition

Applying the Price equation to hierarchical groups requires assigning fitness to lower-level organisms capable of inheritance (Damuth and Heisler 1988; Okasha 2006). This corresponds to Model 1 of multi-level selection (MLS1: (Damuth and Heisler 1988)). In MLS1, the phenotype remains an individual character, and evolutionary change is analyzed through additive summaries of individual traits within groups. Here, the marginal phenotype of the *g*^th^ group (*z_g_*) is defined simply as the mean of the individuals within the group, and the marginal group fitness, *E*[*ɷ* ∣ *g*], is the mean individual fitness within the group (proportional to the group’s total reproductive output, *ɷ_g_* = ∑*_i_*_∈*g*_ *ɷ_i_*). Evolution via multi-level selection is then expressed in terms of two levels of selective covariance and a term for transmission bias

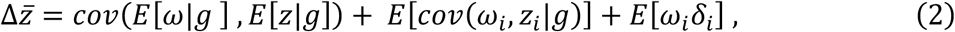

where the first term on the right is the selective covariance attributed to group effects on organism fitness and the second term is the selective covariance attributable to individual fitness independent of any group context. This partition is algebraic and does not, by itself, establish the causal status of either covariance term.

Interpretation of trait–fitness covariance as representing a genuine causal structure requires assumptions about how fitness is generated. In multilevel selection theory, these assumptions take the form of treating additive summaries of group composition (*e.g*., means, sums, frequencies, or linear combinations of individual traits) as sufficient to represent the causal effect of group context on fitness (e.g., (Okasha 2006; Queller 2020)). Accordingly, in the MLS1 approach, once the appropriate additive group descriptor is specified, the detailed configuration of traits within a group should be causally irrelevant to fitness.

This common, but often hidden, assumption can be expressed formally using the concept of screening-off from causal inference. Let *Z_g_* denote an emergent group-level trait that mediates fitness and let *θ_g_* denote an additive group descriptor (*e.g*., the group mean phenotype *z̅_g_* used in Eq. 2). The MLS1 additivity assumption requires that *θ_g_* screens off the dependence of *Z_g_* on the detailed configuration of individual traits *z_i_*:

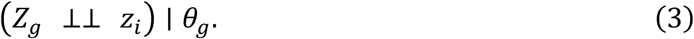

When this conditional independence relation holds, the emergent group mediator *Z_g_* will contain no additional information about fitness beyond what is captured by additive summary of the group composition *θ_g_*; *i.e*., *Z_g_* is causally redundant with respect to fitness. In this case, the causal effect of group context on fitness is fully reducible to the classical partitioning represented by Eq. 2, with the first covariance term on the right interpreted as a genuine group effect on individual fitness (SI Appendix 2).

However, this screening-off condition will not hold in all biological collectives. When there is division of labour, complementarity, threshold effects, or nonlinear synergies, fitness depends on interactions that cannot be reconstructed from any additive summary of group composition. If no additive group descriptor *θ_g_* renders *Z_g_* conditionally independent of individual traits, the effect of group context on fitness is irreducible to independent levels of description. In such cases, individual fitness depends jointly on individual traits and an emergent group configuration, yielding cross-level entanglement of fitness (SI Appendix 2).

The conditional independence relationship defined above is a precise criterion for distinguishing fitness effects that are either reducible or irreducible to identifiable levels of selection. Reducible fitness effects are those for which an appropriate additive group descriptor exists and successfully screens off group configuration. Irreducible fitness effects are those for which no such descriptor exists, leaving individual fitness jointly determined by individual traits and emergent group configuration. The irreducibility criterion delineates a class of fitness effects that cannot be accommodated within additive MLS frameworks; the screening-off criterion introduced here underlies the causal pathway shown in Section 5 and SI Appendix 2. In the next section, this criterion is used to distinguish three qualitatively different kinds of group-level phenomena.

## 4. Three kinds of group effects in multilevel selection

The screening-off criterion introduced in Section 3 is used to classify three different kinds of group-associated fitness effects. Rather than effect size, these are qualitative categories that differ according to whether an additive group descriptor is a causally adequate representation of the group context on fitness. Table 2 summarizes the causal status and reducibility of additive group effects, cross-level by-products, and irreducible effects.

**Table 2.** Causal status and reducibility of group-associated fitness effects.

| Category | Group-level cause | Screening off | Example |
| --- | --- | --- | --- |
| 1. Additive group effect | Yes | Yes (additive descriptor suffices) | Mean toxin-degrading enzyme concentration predicts how ambient toxin levels affect fitness. |
| 2. Cross-level by-product | No | Trivial (conditioning on individual traits eliminates group-level covariance) | Groups of high-fertility individuals produce more offspring (*e.g.*, Fig. 1) |
| 3. Irreducible multi-level effect | Yes | No (configuration dependent) | Division of labour: complementary types must co-occur for high fitness. |

Note: formal conditional-independence expressions are given in Section 3 and SI Appendix 2

### Additive group effects

This is a genuine group effect, where the effect of group context on fitness can be fully captured by an additive predictor that “screens off” the lower-level configuration of the group. Here, conditioning on the appropriate group descriptor renders the configuration of individual traits causally irrelevant to fitness. An example is the collective concentration of a toxin-degrading enzyme. When the enzyme concentration predicts how ambient toxin levels affect fitness, irrespective of the detailed configuration of the population of enzyme produces, then only the mean enzyme concentration is causally relevant to fitness. In such cases, the group covariance term in the Price equation (Eq. 2) can be interpreted as representing a genuine, but additively reducible, group-level effect on fitness. MLS1 contextual analysis (Heisler and Damuth 1987; Damuth and Heisler 1988) also falls within this category when its additivity assumptions are satisfied.

### Cross-level by-products

A limitation of the multi-level Price equation (Eq. 2) is that it makes no distinction between genuine group effects and apparent group-level effects that arise as cross-level by-products of selection acting solely at the individual level (Okasha 2004; 2006). Consider a non-interactive phenotype measured at the group level; say, the mean of individual body mass. Let’s assume that all organisms reproduce at the same time, and that an individual’s reproductive output is positively correlated with its body mass at reproduction. Here, faster growers have higher individual fitness. When total body mass is computed for groups, the distribution will shift towards groups with higher mean body mass as the system evolves (Fig. 1). But, it makes no difference to fitness if an individual finds itself in a group of fast growers or a group of slow growers; the covariance between group mean fitness and group mean phenotype arises despite the absence of any group-level causal mediator. This is an instance of Okasha’s (Okasha 2006) indirect cross-level by-products, the kind Williams (Williams 1966) warned against mistaking for actual group adaptation. In these cases, the apparent group-level covariance disappears once individual-level traits are conditioned upon, indicating that the group term reflects a statistical artifact rather than a causal group effect. Here, screening-off succeeds trivially because there is no emergent group-level cause to screen.

**Fig. 1.**
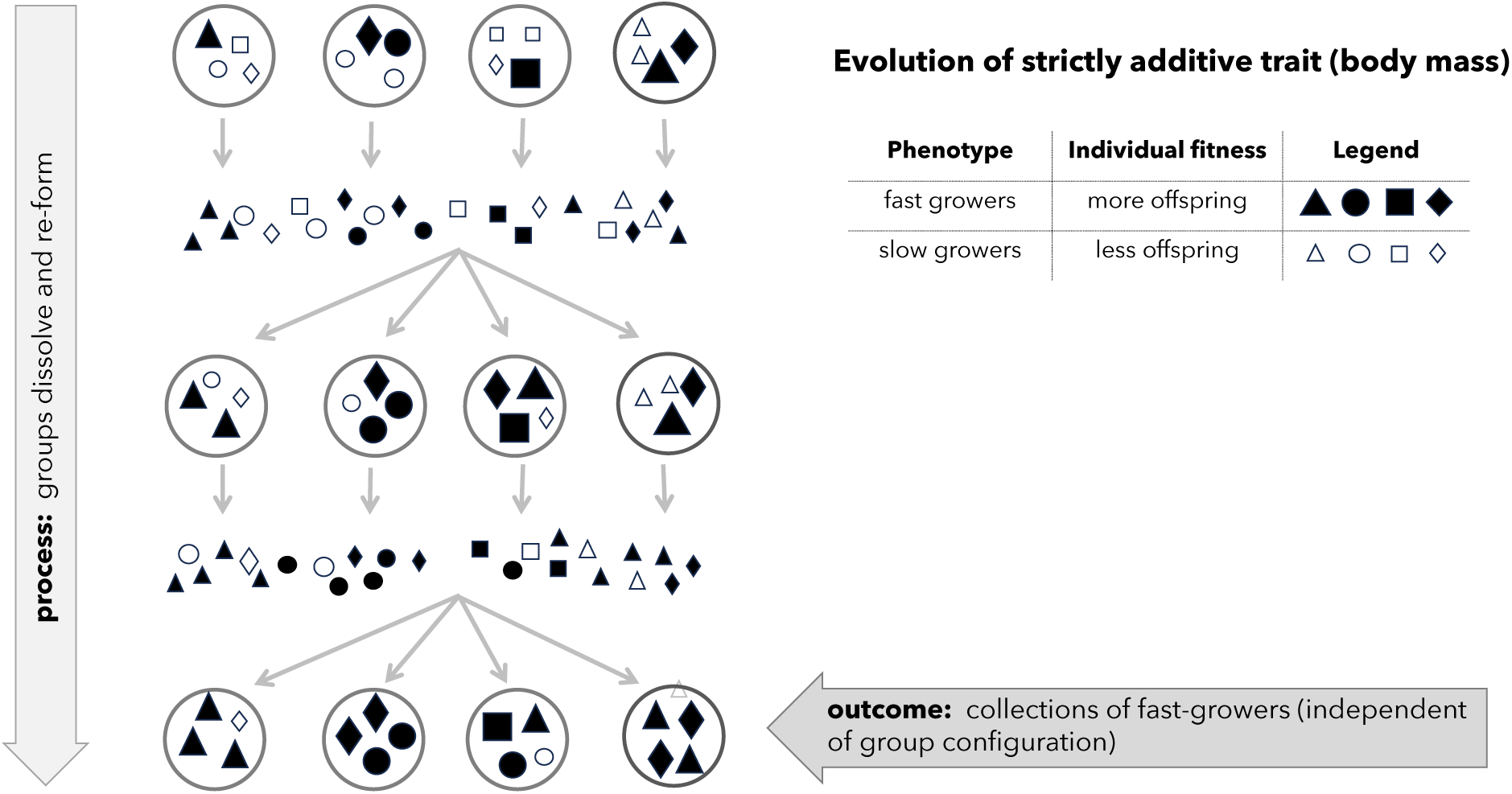
A concept map for the evolution of interspecies interactive genetic effects (IIGEs) by task-group selection. Circles represent multispecies collectives or “communities”, and the symbols (circles, squares, etc.) represent individuals of different species. The dotted lines represent the effect of IIGEs on individual fitness. Each symbol represents an unspecified “dose” of individual organisms belonging to a species. Since collectives dissolve and randomly reform over time they are stochastically “blended” when they re-assemble into a new collective. Because individuals benefiting from fitness enhancing IIGEs produce more offspring, collectives that contain more positively interacting individuals will, on average, produce more offspring. In this way the among-group distribution of IIGEs evolves toward greater representation of mutualistic interactions within groups. Note that this model works just as well when symbols represent different kinds of organisms within a species, with mutualistic IGEs evolving within a single species. In the context of the Price equation for MLS1, such mutualistic interactions are selected effects, despite the absence of a collective-level inheritance mechanism.

### Irreducible multi-level effects

A unique kind of group effect will arise when group context affects fitness through emergent, configuration-dependent interactions that cannot be represented by any additive group descriptor. A generic model of task group selection (Griesemer 2014) is used to illustrate the evolution of collective-level interactions without group-level heredity (Fig. 2). Note that the symbols in Figure 2 (circles, squares, etc.) represent individuals of different species, and the dotted lines represent IIGEs, but this model also works when symbols represent different kinds of organisms within a species and the dotted lines represent IGEs. Here, individuals having fitness enhancing interactions within their group produce more offspring. Because groups that contain more positively interacting individuals will, on average, contribute more lower-level offspring, the among-group distribution of interactive effects (IGEs or IIGEs) evolves in response to their effect on individual reproduction. Critically, when no additive summary can screen off the causal influence of a group configuration, individual level fitness will depend jointly on individual traits and emergent group structure. The resulting cross-level covariance reflects *irreducible causal entanglement* across levels rather than an additive decomposition of effects. Such irreducible group effects differ fundamentally from the additive group effects and cross-level by-products described above; this is a genuine group causal effect that cannot be decomposed into additive, gene- or individual-level contributions without loss of causal information (SI Appendices 2-4). Importantly, these effects qualify as selected effects even in the absence of a mechanism for group-level inheritance.

**Fig 2.**
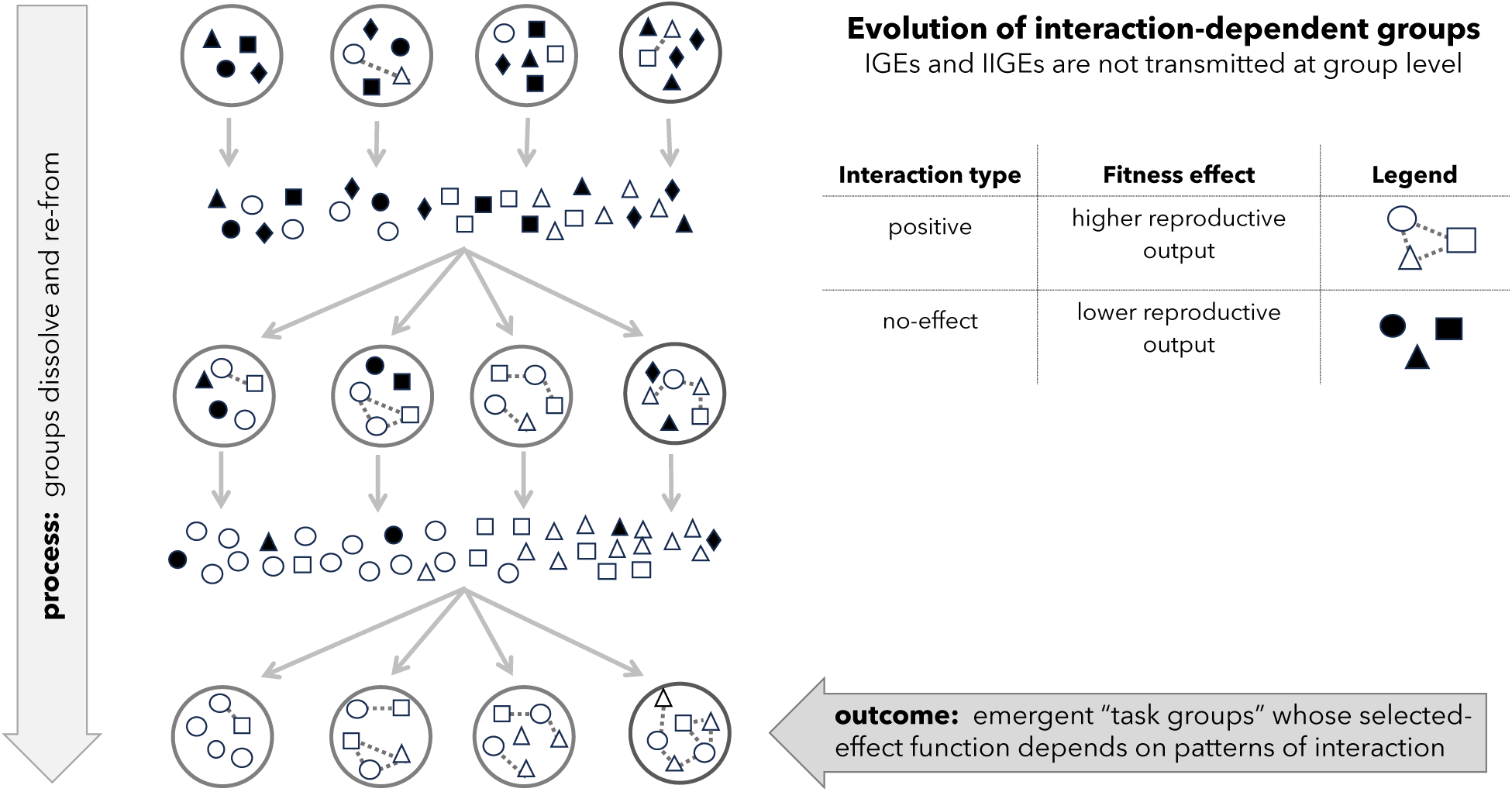
A concept map for the evolution of a *non-interactive* phenotype measured at the group level. Circles represent multispecies collectives, and the symbols (circles, squares, etc.) represent individuals of different species. There are no interactive effects on individual fitness. All organisms reproduce at the same time, and an individual’s reproductive output is simply a positive function of its own body mass at reproduction. Hence, it makes no difference to individual fitness if an individual finds itself in a group of fast growers or a group of slow growers. Collectives dissolve and are stochastically “blended” when they re-assemble into new collectives. When total body mass is computed for groups, the distribution shifts towards higher group values as the system evolves because there is a statistical covariance between aggregate group body mass and aggregate individual fitness. This example illustrates that group-level statistical covariance is not equivalent to group-level selective covariance (*c.f.* Fig 1.).

### Boundary cases

A recent gene-centric approach explicitly incorporates complex interactive effects, including non-transmitted and non-additive social interactions, by representing them as additive contributions to lower-level fitness (Queller 2020). While this greatly expands the range of interactions admissible within a selective explanation, it does so by imposing a constraint: all causal effects must be expressible as additive predictors attributable to genes. As a result, even this interaction-friendly gene’s-eye framework presupposes screening-off by additive descriptors. Hence, this gene’s-eye-view systematically excludes causal structures in which fitness depends irreducibly on group configuration. Hull’s interactor–replicator framework (Hull 1980) similarly relaxes heredity requirements at higher levels, but remains silent on the causal architecture of interactors and does not distinguish cases in which their fitness effects are additively decomposable from those in which they depend irreducibly on group configuration. Both of these are boundary cases because they accommodate interaction effects only insofar as those effects can be expressed as additive predictors that screen off group configuration. Understanding how cross-level interactions can generate emergent fitness that cannot be screened off requires a causal modelling approach.

## 5. Synaptive MLS: Cross-Level Causality and the Evolution of Interactive Traits

### 5.1. Structural origin of irreducible cross-level covariance

Natural selection, as classically understood, requires a covariance between fitness and heritable trait transmission (Eq. 1). Yet, selective covariance can arise at higher levels of organization without inheritance mechanisms at those levels. When fitness is shaped by collective traits emerging from inter-individual interactions (*e.g.*, Fig. 2), trait–fitness covariance can become irreducible across levels; that is, the causal structure of fitness cannot be represented by the additive, level-specific, contributions given by the MLS1 Price decomposition (Eq. 2). Bijma’s (Bijma 2020) Price-equation treatment of IGEs shows how interactions generate trait–fitness covariance, but because those effects are expressed entirely through additive predictors they are fully reducible. In contrast, the case considered here involves cross-level entanglement, where no additive set of predictors can screen off (*i.e*., render conditionally independent) how emergent non-linear interactions affect the fitness of lower-level individuals. Level-specific causal attribution will be inappropriate and inadequate in such multi-level systems (Lloyd 2005). This scenario is analyzed formally in SI Appendix 2, which shows that irreducible selective covariance arises when an emergent group-level mediator cannot be rendered conditionally independent of individual traits given any additive group-level predictor.

A causal graph (Fig. 3) shows how collective mediators of non-additive fitness (e.g., IGEs and IIGEs) can entail causal entanglement, with individual fitness being shaped by emergent group-level interactions. Figure 3 shows how individual-level traits (*z_i_*, *z_j_*) in group *g* can influence fitness through direct and emergent pathways. Traits of interacting individuals jointly determine the interaction term, *I*(*z_i_*, *z_j_*), which contributes to the construction of a group-level emergent trait *Z_g_*, such as task specialization or social coordination. This trait, in turn, influences the fitness of the interacting individuals (*ɷ_i_*, *ɷ_j_*) in group *g*. Because *Z_g_* reflects the non-linear integration of multiple contributors, its effects cannot be reduced to the additive partitioning assumed by MLS1 (SI Appendices 2-4). Emergent mediation of fitness can arise in diverse systems, including microbial and insect divisions of labour, human social interactions, early multicellular aggregates, and holobionts.

**Fig 3.**
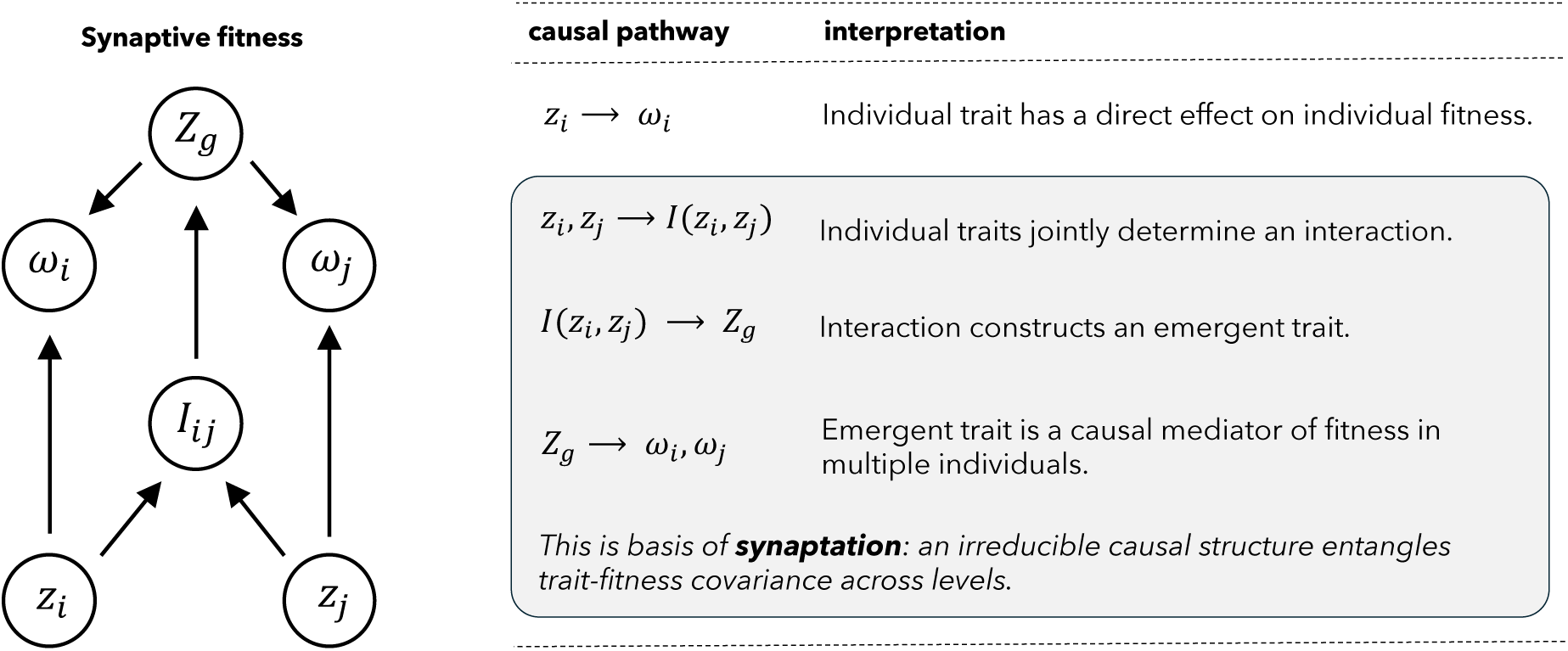
The causal structure of Synaptive MLS1 and irreducible cross-level dependencies between trait and fitness. The trait value in the *i^t^*^h^ individual (*z*_*iG*_) and *j^t^*^h^ individual (*z*_*jG*_) in the *G*^th^ group affect their individual fitness (*ɷ*_*iG*_ and *ɷ*_*jG*_ respectively) by (i) a direct pathway and (ii) indirectly via an emergent mediator. The indirect pathway is mediated by an interaction (*I*_*ijG*_) whereby the individual traits contribute to the construction of a group-level emergent trait (*Z*_*G*_), which also influences the fitness of the interacting individuals (*ɷ*_*iG*_, *ɷ*_*jG*_). The dependence of fitness on an emergent, interaction-driven, trait means that selective covariance is causally entangled across levels.

### 5.2 Synaptation: definition, taxonomy, and continuum

A new term is needed to represent the distinct selective origin of traits produced by task-group selection because such traits: (*i*) serve a novel selected-effect function, and (*ii*) arise through irreducible cross-level causation mediated by distinct selected effects at lower-levels. I propose the term *synaptation* for this interactive complexity (Table 3). The interactive component of individual *fitness* (*aptus*) is gained only when an individual *acts with* (*syn*) other individuals of the relevant kind; *i.e*., is *synaptus*, within its group (visualized in Fig. 2). As a selected-effect trait, a synaptation is defined by irreducible cross-level selective covariance (SI Appendix 2). Synaptation extends recent realist accounts that emphasize how non-aggregative traits are evidence of genuine collective function (e.g., (Bourrat 2021)) by identifying the irreducible cross-level causal structure that cannot be eliminated by additive group-level description. This poses a problem for the current trait taxonomy (Table 1) because the evolutionary origin of a synaptation cannot be localized to an individual phenotype nor a heritable group-level trait, yet it integrates multiple adaptive or exaptive selected effects into a single higher-level function.

**Table 3.** An expanded taxonomy of fitness-related traits.

| Character | Status | Co-option | Historical Function | Selective Causation | Fitness of entity | Semantic Category |
| --- | --- | --- | --- | --- | --- | --- |
| Aptation | A trait that currently enhances fitness, making no claims about its evolutionary origin. A super-category that includes adaptation, exaptation and synaptation. | unspecified | unspecified | unspecified | unspecified | unspecified |
| Nonaptation | A trait that currently does not affect fitness, making no claims about its evolutionary origin. | no | unspecified | not at present | not at present | neutral |
| Adaptation | A trait that currently enhances fitness *and* was built by natural selection for the use it currently serves. | no | continuous | level-specific | directly enhanced | function |
| Exaptation<sup>A</sup><br>(from adaptation) | A trait that was built by natural selection for a particular function (i.e., adaptive at the time of its origin) *and was coopted* for a new use that currently enhances fitness. | yes | previously adaptive | level-specific | directly enhanced | effect |
| Exaptation<sup>N</sup><br>(from nonaptation) | A trait that was previously nonaptive *and was coopted* for a new use that currently enhances fitness. | yes | previously nonaptive | level-specific | directly enhanced | effect |
| Synaptation | An emergent collective trait that was built for its current use by multi-level selection, but which integrates individual traits and behaviours that can *simultaneously have other functions and effects*. | yes: dual-aspect<sup>1</sup> | emergent<sup>1</sup> | irreducible across levels | interaction mediated<sup>2</sup> | coordinated function and cross-level effects |

<sup>1</sup> Synaptation involves dual-aspect recruitment, where the lower-level traits preserve their original selected functions while becoming integrated into an emergent function at a higher-level where they contribute to a distinct selected-effect. <sup>2</sup> Reciprocal interactions within a heterogenous collective mediate individual fitness, generating cross-level causal entanglement.

Synaptation will generate a non-zero MLS1 between-group covariance. However, this covariance will be *irreducible* because no additive group predictor (e.g., a group mean) can screen off the configuration-dependent interactions that construct the emergent mediator *Zg* (SI Appendix 2). When the MLS1 reducibility assumption fails, the resulting between-group covariance reflects genuine cross-level entanglement rather than a canonical additive group effect. In these cases, the classical MLS1 formulation misattributes configuration-dependent fitness interactions to an additive group mean (SI Appendices 3–4). Synaptive MLS makes the non-decomposable causal architecture explicit, showing how fitness is mediated by emergent, cross-level interactions rather than by traits at any single level (Fig. 3; SI Appendix 2).

The historical genesis of a synaptative trait reflects a multilevel process of evolutionary cooption that is neither strictly adaptive nor strictly exaptive in the classical sense. Group-level interactions shaped by synaptive MLS will depend on individual-level traits that may retain their original adaptive functions (*i.e*., functions tied to their lower-level selective origin), or may retain their derived roles as exaptations^A^ or exaptations^N^, or may even have no current fitness value at the individual level. In some cases, a synaptation might even incorporate traits that are mildly deleterious (inapt) from the individual-level perspective. Synaptive MLS thus offers a mechanism by which traits originally shaped for disparate roles, or are even non-functional traits, can scaffold the emergence of novel collective functions. In this context, the term exaptation is inadequate: synaptations are not fully formed traits re-purposed for some new role in the same unit of selection, but emergent levels of functional organization that depend on traits that simultaneously serve other functions and effects at lower levels (Fig. 3). Nor are synaptations pure adaptations in the classic sense, as they originate by co-opting and re-purposing a set of lower-level traits. Synaptations integrate heterogeneous evolutionary histories into coherent functionally selected group-level traits. Taking these considerations together with the distinct causal architecture of synaptations motivates the expanded fitness taxonomy developed in Table 3.

Synaptation represents more than a new category in the taxonomy of fitness. Together with adaptation, they define the endpoints of a continuum of selected-effect complexity (Table 4). At the adaptive end, traits such as the vertebrate eye illustrate systems in which the fitness effects of heterogeneous components are fully assimilated through shared development and a single hereditary lineage. At the synaptive end, holobionts and other multi-lineage collectives illustrate systems in which heterogeneous fitness contributions are mediated by cross-level interactions. Adaptations and synaptations are both historical selected-effect functions, but they differ in causal architecture: adaptations arise from additively decomposable fitness effects optimized within one lineage, whereas synaptations arise from irreducible cross-level dependencies distributed across lineages. Although their diagnostic covariance structures remain binary (additively reducible for adaptations, and irreducible for synaptations) biological systems can blend these causal architectures in generating fitness. Thus, adaptation and synaptation define a continuum in how selection integrates causal architectures across levels (Table 4). The dual-aspect origin of the synaptive traits explains how lower-level components retain their ancestral causal roles, or repurposed effects, while contributing to the origin of new emergent traits with higher-level functions.

**Table 4.** Synaptation and adaptation define the endpoints of a continuum of selected-effect complexity.

|  | **Adaptation** | **Intermediary** | **Synaptation** |
| --- | --- | --- | --- |
| Causal structure | Additive selective covariance reducible to a single level (individual or group) | Partially coupled levels with mixed causal architectures | Cross-level, irreducible selective covariance |
| Inheritance | Single lineage with unified heredity | Partially distributed heredity | Distributed across autonomous but interacting hereditary systems |
| Fitness mediation | Homogeneous, internal processes | Mixed, with some mediation via division of labor or symbiosis | Heterogeneous, interaction-dependent processes mediated by emergent collective properties |
| Fitness alignment | Intrinsic mechanisms suppress conflict (e.g., germline control, fair meiosis, apoptosis, epigenetics) | Mixed, with partial subordination of conflicts | Extrinsic mechanisms coordinate compatible but diverse fitness consequences that emerge through cross-level interaction (e.g., ecological feedbacks) |
| Origin of complexity | Natural selection integrates heterogeneous traits into a unified system where fitness is optimized by a single evolutionary lineage | Natural selection blends integration and coordination of heterogeneous traits in partially coupled systems. | Natural selection coordinates heterogeneous traits that serve other selected-effect functions across multiple levels and evolutionary lineages |

### 6. Example of synaptation: the vampire bat + microbiome holobiont

The extant species of “vampire bats” (subfamily Desmodontinae) are obligate blood feeders. Blood is an extremely challenging source of nutrition because it contains very little carbohydrates, almost no vitamins, and it entails potentially high exposure to blood-borne pathogens (Breidenstein 1982; Riskin and Carter 2023). To survive solely on blood a vampire bat depends on an impressive spectrum of anatomical and physiological traits (Schmidt 2018). The physiological traits include saliva-mediated anticoagulation, specialized interferon responses to pathogens, and a physiologically integrated gut microbiome (Riskin and Carter 2023; Zepeda Mendoza et al. 2018). The classic evolutionary interpretation holds that all the vampire bat’s unique traits are individual-level adaptations, each serving *the fitness of the individual bat* (*e.g.*, (Phillips and Baker 2015; Blumer et al. 2022; Ribeiro et al. 2022)). However, the physiological capacities encoded in a vampire bat’s genome is insufficient to support its survival as an obligate sanguivore; the lifestyle cannot be sustained without the functions and effects also encoded within its gut microbial community ((Suárez and Triviño 2020) and references therein)).

Vampire bats form colonies ranging from a few individuals to hundreds and are socially structured around matrilines where some female offspring remain in the colony along with a few temporary adult males. Relatedness within colonies remains low due to the frequent immigration of unrelated females, random male dispersal, and expulsion of older males (Wilkinson 1985). In *Desmodus rotundus*, these factors result in population structure with lower-than-expected within-group relatedness. In addition, vampire bats rely heavily on social behaviors: individuals often regurgitate blood to feed both kin and non-kin, mediated by social bonds such as allogrooming and reciprocal cooperation (Wilkinson 1984; Carter and Wilkinson 2013; 2015; Carter et al. 2017). These interactions facilitate the formation of a “social microbiome” where mouth-to-mouth feeding and grooming lead to convergence of gut microbial communities across social networks (Yarlagadda et al. 2021). Given widespread bat dispersal among colonies and the lateral transmission of microbes, neither colonies nor holobionts qualify as a canonical unit of selection, despite functioning as coherent biological interactors (Lloyd 2018). Yet, a vampire bat’s fitness depends on both conspecific social partners (via IGEs) and microbial symbionts (via IIGEs). The evolution of sanguivory likely required some form of task-group selection.

Based on a community-wide analysis of adaptive genomic signatures, Zepeda Mendoza et al. (Zepeda Mendoza et al. 2018) identified fitness-affecting dependencies between the genetic elements of vampire bats and their gut microbiomes. These constitute a defined set of putative IIGEs that support a collective capacity for sanguivory. For example, genes linked to starvation or low-nutrient responses (*e.g.*, *LAMTOR5*) exhibit signs of positive selection in the vampire bat genome and are mirrored by enrichment of complimentary functions in the gut microbiome (*e.g*., *RelA/SpoT* family and guanosine pentaphosphate pathways). Other areas of complementary specialization include blood viscosity/coagulation, management of nitrogen waste & blood osmotic pressure, and assimilation of lipid, glucose & iron. Suárez and Triviño (Suárez and Triviño 2020) describe the difficulty with treating these complementary specializations as adaptations that belong exclusively to either the vampire bat or its gut microbes. They proposed viewing this multispecies system as a *holobiont*, a collective entity exhibiting adaptations for sanguivory. While this holobiont does indeed represent a complete informational entity in so far that it would contain the IIGEs required for the “task” of sangivory, it does not address how multi-level selective covariance explains the origin and maintenance of holobiont level fitness when the holobiont does not sustain heredity.

While obligate sanguivory is best understood as a property of the holobiont (*cf*. *Z_g_* in Fig. 3), it is not a unit of selection according to Lewontin’s criteria (Zepeda Mendoza et al. 2018; Suárez and Triviño 2020). Here, the concept of synaptation, arising from task-group selection, is helpful. The collective task of modulating blood coagulation and viscosity during feeding (*cf*. *I*(*z_i_*, *z_j_*) in Fig. 3) serves as an example. Vampire bats facilitate feeding by secreting a salivary enzyme, encoded by *ENTPD1*, that modulates wound coagulation (*cf*. *z_i_* in Fig. 3). Originally, the *ENTPD1* enzyme inhibited blood clotting within the circulatory system, making its feeding-related role an *exaptation* of its ancestral function. The exaptive shift to a role in feeding occurred in an insectivorous ancestor, likely facilitated by regulatory changes arising from adaptive or neutral mutations affecting salivary gene expression. Thus, secondary exaptation of the *ENTPD1* enzyme for sanguivory required complementary functions within the gut microbiome (*cf*. *z*_*j*_ in Fig. 3), which provides the microbial enzymes that degrade host coagulation enhancers and reduce coagulation factor synthesis (Zepeda Mendoza et al. 2018). Any irreducible selective covariance for these complementary enzymatic functions during the ∼4 million-year origin and divergence of vampire bats from insectivorous ancestors makes the collective task of modulating blood coagulation a *synaptation*. This synaptation is built from a variety of traits that include exaptive enzymatic effects within individual vampire bats and bacteria, adaptive or neutral changes in bat gene expression, and secondary adaptation of proteins that fine-tune the system (*i.e*., the adaptive signatures detected by Zepeda Mendoza et al. (Zepeda Mendoza et al. 2018). Management of blood coagulation is a complex relational trait that is *simultaneously* a synaptation at the level of the holobiont and a mix of exaptations^N^, exaptations^A^, and adaptations at the level of individual organisms.

## 7. Discussion

Griesemer (Griesemer 2014) introduced the concept of a *task group* to clarify how heterogenous entities could perform functions that no single individual could achieve, often without group-level heredity. In *task-group selection* the fate of individuals is *coordinated rather than shared*, typically through division of specialized activities. Because the members of a task group play complementary roles, they experience diverse fitness consequences. This differs from Wilson’s (Wilson 1980) *trait group selection* where, by contrast, the members of a group share a common trait that links them to a shared fate. The concept of synaptive MLS extends task-group selection by describing the evolutionary origin of hybrid selection products that emerge through irreducible cross-level covariance. Thus, *synaptation* gives a name to a selection product with heterogenous fitness consequences, whereas *synaptive MLS* describes the process by which it originated.

The extended fitness taxonomy presented here situates selection products as the endpoints of a continuous spectrum of causal integration. At one endpoint, adaptations evolve through lineage-bound selection, where fitness is fully decomposed into additive effects transmittable via a shared hereditary system. At the other endpoint, synaptations evolve when fitness is non-decomposable and distributed across interacting entities with distinct hereditary mechanisms and evolutionary histories. This spectrum reframes evolution of complexity as a process of causal integration that can progress in different dimensions, sometimes leading to complex traits with blended origins. For example, Roughgarden’s (Roughgarden 2023) theoretical model for holobiont evolution blends community-level partner evolution (MLS1) with adaptive evolution of host-level recruitment mechanisms, showing how feedback between them generates strong holobiont-level selection without holobiont-level heredity. An empirical example of blended origins is the synaptive thermoregulation of brood temperature in honeybees. The fitness of male drones and the queen depend on maintaining brood temperature in a narrow range through coordinated, non-linear integration of heterogeneous sub-traits (fanning, clustering, moisture regulation, colony architecture). Kin selection ensures that the fitness benefit of colony-level thermoregulation accrues differently in the adaptive evolution of genes originating from the queen versus those originating from male drones. Nonetheless, the emergent functional coherence of brood thermoregulation leads to causal entanglement of their fitness. Framing adaptation and synaptation as endpoints of a causal continuum bridges the classical boundary between inheritance-based and cooperative-emergent evolution, showing that natural selection can operate via both the assimilation and the coordination of fitness effects.

Holobionts, task groups, and cultural collectives share a common feature: functionally coordinated division of labour. For this reason synaptation applies broadly, from conspecific collectives and multi-species communities (Queller 2020; Lean et al. 2022; Lean and Jones 2023; Doolittle 2024a; Papale and Doolittle 2024) to cultural systems (Griesemer 2014; Doolittle 2024b). In multi-species systems, task groups are hybrid entities composed of individuals from distinct lineages. For instance, metabolic cooperation of bacterial species in the human oral microbiome is mediated by non-homologous IIGEs that generate non-additive, cross-level fitness effects (Elias and Banin 2012). Single-species systems can also evolve fitness entanglement through division of labor. In genetically chimeric collectives such as the slug and fruiting body stages of the social amoeba *Dictyostelium purpureum* (Strassmann et al. 2000), heterogeneous genotypes are integrated through irreducible covariance between individual and collective fitness. Finally, cultural division of labor is scaffolded by socially transmitted traits: heterogenous cultural resources flow both horizontally and vertically among members, supporting hybrid social groups with emergent function that can feed back on the reproductive success of individual group members (Griesemer 2014). Synaptive MLS offers a common causal architecture for cases such as these, where individuals in heterogeneous groups evolve coordinated functions while operating on distinct fitness landscapes.

At this point, the question is no longer whether these systems exist, but how their selected effects originate across distinct fitness landscapes. Queller (Queller 2014) used the metaphor of multiple adaptive landscapes to illustrate how multi-species evolution is constrained by selective conflict. Extending this metaphor, task-group selection can be likened to teams of mountain climbers navigating separate landscapes while tethered by ropes, the IIGEs, which link their evolutionary outcomes. Though each team member ascends its own landscape, their trajectories are interdependent: success or failure is propagated through their synaptive genetic linkages, forming the multi-species “survival tool-kit” (Fig. 4A). Models that treat IGEs or IIGEs as components of a shared phenotype show that within-group selection can increase population mean fitness via their additive effects (Queller 2014; De Lisle et al. 2022). Yet these same genetic interdependencies impose constraints in which selection increases fitness only up to the point where fitness conflicts stall further gains (Queller 2014). Synaptive MLS extends such single-level models by adding an evolutionary mechanism through which certain collectives, with their uniquely optimized interactions and minimized conflicts, can outperform other collectives over evolutionary time (Fig. 2).

**Fig 4.**
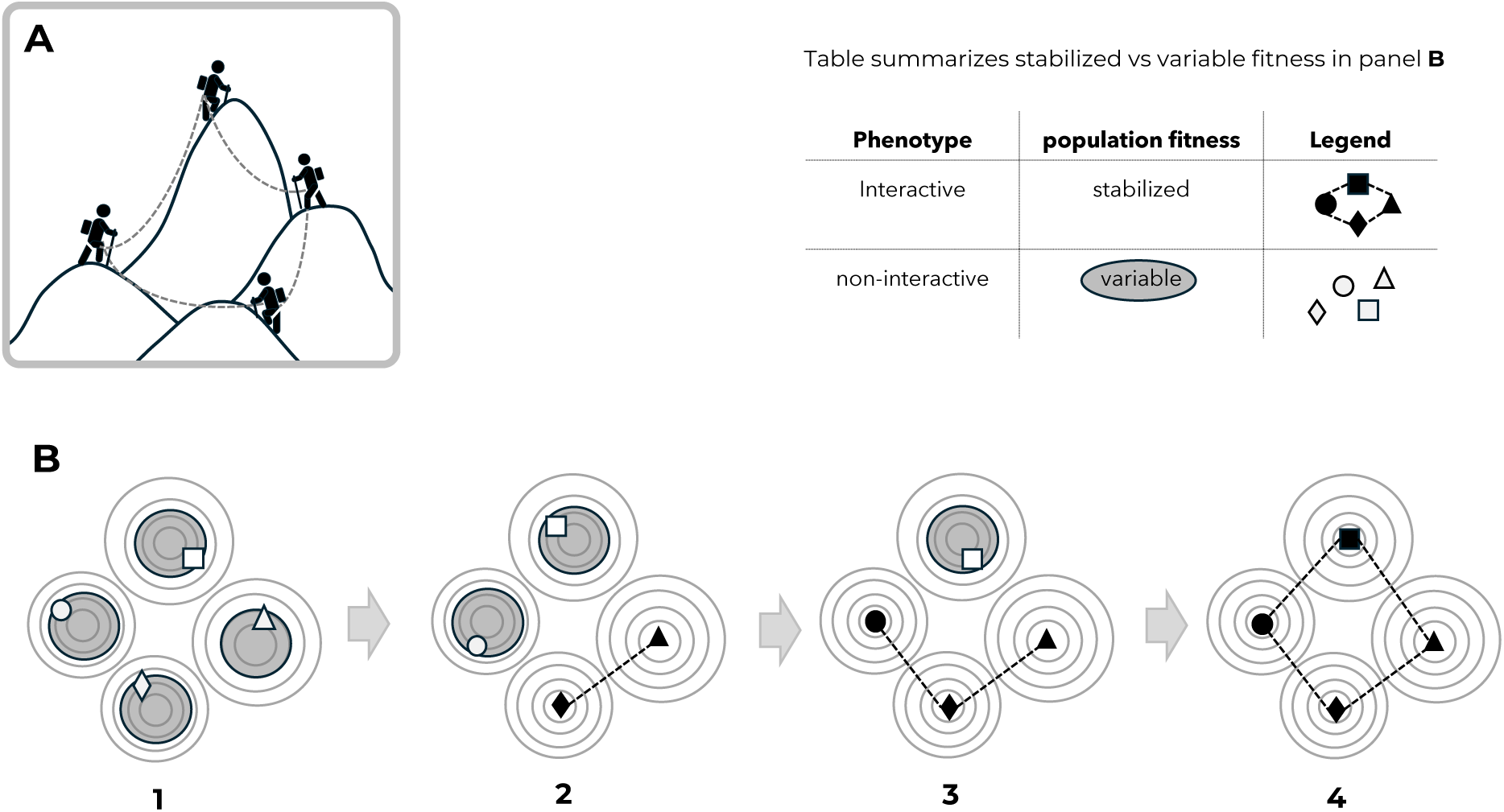
Evolution of interactions on a multi-species fitness landscape. The adaptive landscape metaphor is extended to multi-species interactions arising from inter-specific interactive genetic effects (IIGEs) on fitness. Each species ascends its own landscape, but their successes or failures are distributed through their synaptive genetic linkages. Collectives that interact in this way are like teams of mountain climbers navigating separate landscapes while tethered by ropes, the IIGEs, which link their evolutionary outcomes (**A**). Synaptive MLS1 is an evolutionary mechanism through which certain collectives (“teams”), with their uniquely optimized interactions and minimized conflicts, can outperform others over evolutionary time. Rather than precisely occupying a fitness peak, natural populations experience a dynamic state of “shifting balance” due to the interaction of selection, drift, and other forces (**B**). Because IGEs and IIGEs can alter heritability and reshape fitness landscapes, their evolution by task-group selection could transform within-species evolutionary dynamics and potentially stabilize outcomes. In this way task-group selection could lead to evolutionary transitions in trait coordination among species, thereby growing cross-level entanglement of non-linear fitness effects.

The metaphor of populations steadily climbing fitness peaks is an oversimplification. Fitness does not always increase; dominance, epistasis, frequency-dependent selection, and drift can cause fitness to decline (Crespi 2000; Kvitek and Sherlock 2011; Brady et al. 2019). A key question is whether the recruitment of IGEs or IIGEs contributes not to maximization, but to stabilization of fitness. In figure 4B four species initially occupy regions of a landscape (not peaks) because stabilization is prevented by ongoing drift and selection (*e.g.*, (Jones et al. 2017)). Because interactive genetic effects can alter within-species heritability (Wolf et al. 1998; Bailey and Desjonquères 2022) and reshape fitness landscapes (McGlothlin et al. 2010; Bijma 2011), task-group selection could transform within-species evolutionary dynamics and potentially stabilize collective outcomes (Fig. 4B). Niche construction can also increase heritability and reshape fitness landscapes, thereby extending the influence of IGEs and IIGEs across generations (Fogarty and Wade 2022). Such transgenerational effects, often termed *ecological inheritance* (Odling-Smee 1988; Rossiter 1996; Laland et al. 2015), could become the target of task-group selection thereby reducing the sensitivity of niche-constructing task-groups to random assembly. Because these effects are not part of classical quantitative genetic theory (Fogarty and Wade 2022), the role of task-group selection in modifying heritability and fitness landscapes represents an promising direction for future work on understanding evolved complexity.

Emerging notions of relational health (Pitt and Gunn 2024) presuppose multispecies coordination and stabilization despite divergent fitness consequences. Synaptive MLS identifies the origins of such coordinating mechanisms, connecting evolutionary causation to relational concepts of fitness and health. Within this framework, eubiosis represents a synaptive equilibrium where cross-species or cross-individual interactions stabilize irreducible function. Dysbiosis arises when synaptive functions break down, and rebiosis is their restoration through reassembly or reactivation of interactive causal structures. The human social microbiome exemplifies this process: mutualistic microbes transmitted through social contact contribute to immune regulation (Sarkar et al. 2024), consistent with the “Old Friends” hypothesis that coevolved microbial interactions calibrate immune tolerance (Rook 2023). Viewed synaptively, human health becomes an emergent selected effect arising from persistent coordination among heterogeneous biological and social partners. Disruption of these causal linkages (dysbiosis) can propagate dysfunction across levels, whereas rebiosis restores functional integration (Verburgt et al. 2023). In this way, synaptive MLS serves as a general framework for understanding eubiosis, dysbiosis, and rebiosis as eco-evolutionary processes that maintain or recover cross-level causal coherence in health-related systems.

The source of variation from which new complexity evolves is also broadened by the concept of synaptation. Conventional mutation, recombination, and gene duplication generate variation randomly with respect to fitness (e.g., Kliebenstein 2008; Magadum et al. 2013), and synaptation extends this by revealing how selected-effect traits at one level can become the pool of random variation required for higher-level evolutionary innovation. Lower-level adaptations and exaptations thus provide the variation that scaffolds the emergence of new functional hierarchies. Because selective covariance in synaptive systems is irreducible across levels, its historical genesis cannot be captured by a single unit of selection (Lloyd and Gould 1999; Lloyd 2005; 2018). Synaptation removes the pressure to identify the primary unit of selection. By showing how collectives can evolve novel selected-effect functions while their constituent parts retain their distinct adaptive or exaptive roles, synaptive MLS offers a general mechanism for how selection builds complexity from coordination among heterogenous entities.

Together with recent work on persistence-based selection (e.g., (Bourrat 2021; Neto and Doolittle 2023; Papale and Doolittle 2024; Doolittle 2025)), synaptive MLS supports an expanded view of how natural selection shapes function and complexity beyond inheritance-bound systems. Both perspectives reject the idea that complex adaptations must arise solely through upward causal flows from lower-level selection (e.g. (Dawkins 1976; Maynard Smith 2002)), and instead emphasize that emergent functions can evolve through selective coordination of heterogeneous entities. Synaptive MLS make this clear by describing how irreducible cross-level covariance generates functional coherence across distinct evolutionary lineages. It also offers a framework for interpreting complex multi-species phenomena, such as ocean metabolism (Saito et al. 2024) and hologenome evolution (Rosenberg and Zilber-Rosenberg 2018), as outcomes of selection acting through cross-lineage mediation of non-homologous IIGEs. Such synaptive complexity is heterogeneous by construction, serving a selected function at one level while being composed of components that are simultaneously adaptations or exaptations for lower-level units of selection. Together, synaptive- and persistence-based approaches advance evolutionary theory toward explaining how natural selection contributes to life’s complexity beyond inheritance-bound systems.

## Acknowledgments

I am grateful for Letitia Meynell’s and François Papale’s helpful discussion of the relationship between adaptation and synaptation. Discussions with Ed Susko on the topic of irreducible cross-level covariance in causal models were especially valuable. Ed Susko also pointed out the value of a minimal numerical example of irreducible synaptive covariance, for which I am very grateful. I thank Ford Doolittle, Letitia Meynell and François Papale for highly formative discussions and their insightful comments on the manuscript. The paper was written with the financial support from the New Frontiers in Research Fund (NFRFE-2019-00703) and NSERC Discovery Grant(s) (RGPIN-2020-04109; RGPIN-2025-03979).

