## Supplemental Appendices 1-5 for "Synaptation − a missing concept in the science of evolved complexity"

Joseph P. Bielawski

Corresponding author:

*Joseph P. Bielawski, Department of Biology, Dalhousie University, Halifax, NS, Canada.*

**This PDF file includes:**

- **SI Notation**
- **SI Appendix 1:** Natural Selection Requires Additive Effects at One Level.
- **SI Appendix 2:** Irreducible Cross-Level Covariance and the Limits of MLS1
- **SI Appendix 3:** A Minimal Numerical Example of Irreducible Synaptive Covariance
- **SI Appendix 4:** A Biologically Intuitive Example: Division-of-Labour Synaptation
- **SI Appendix 5:** A Diagnostic for Synaptation (Irreducible Cross-Level Selective Covariance)
- **SI References**

#### SI Table of Contents

| Section | Title | Page |
|---|---|---|
| **SI Notation** | **Notation Used Throughout SI……………………………………………..** | **3** |
| **SI Appendix 1** | **Natural Selection Requires Additive effects on fitness at one level……….** | **4** |
| **SI Appendix 2** | **Irreducible Cross-Level Covariance and the Limits of MLS1……………..** | **6** |
| 2.1 | Purpose…………………………………………………………………... | **6** |
| 2.2 | Causal Model and Notation…………………………………………….... | **6** |

#### Notation Used Throughout SI

**Indices and structure**

| | |
|---|---|
| $i, g$ | indices for individuals and groups |
| $n_g$ | number of individuals in group $g$ |

**Traits and phenotypes**

| | |
|---|---|
| $z_i$ | trait (phenotype) of individual $i$ |
| $\bar{z}, \bar{z}_g$ | mean trait across all individuals; mean trait in group $g$ |
| $z_{-i}$ | trait vector of all individuals in group $g$except $i$ |
| $z_i^o$ | mean offspring trait of individual $i$ |
| $\delta_i$ | parent–offspring trait difference |

**Fitness**

| | |
|---|---|
| $w_i, \omega$ | individual fitness |
| $\bar{w}_g = E[w_i \mid g]$ | mean fitness in group $g$ |

**Price equation and quantitative genetics**

| | |
|---|---|
| $\Delta\bar{z}$ | change in mean trait across generations |
| $\mathrm{Cov}(\cdot,\cdot), \mathrm{Cov}_g(\cdot,\cdot)$ | covariance across individuals; covariance across groups |
| $E[\cdot]$ | expectation operator |
| $S$ | selection differential, $\mathrm{Cov}(\omega, z)$ |
| $V_A, V_P$ | additive genetic variance; phenotypic variance |

**Additive group descriptors (screening variables)**

| | |
|---|---|
| $\theta_g$ | generic additive group-level descriptor (mean, sum, frequency, or linear combination); candidate screening variable |

**Emergent and configuration-dependent variables**

| | |
|---|---|
| $Z_g$ | emergent group-level mediator arising from configuration-dependent interactions |
| $Z_g^{(2)}$ | second moment of trait values within group $g$ |
| $A_g$ | vector of additive summaries of group composition (means, sums, frequencies, additive moments). |
| $C_g$ | vector of configuration-dependent group descriptors |
| $E_g$ | emergent group-level mediator used in the diagnostic framework |

**Causal and functional notation**

| | |
|---|---|
| $\phi(\cdot)$ | Causal fitness function |

**Regression and diagnostic framework (SI Appendix 5)**

| | |
|---|---|
| $\alpha, \beta_{\text{ind}}, \gamma_j, \delta$ | regression coefficients (intercept; individual effect; additive group effects; emergent mediator) |
| $\varepsilon_i$ | regression error term |
| $H_0, H_A$ | null and alternative hypotheses |

**Conditional independence and testing**

| | |
|---|---|
| $(X \perp\!\!\!\perp Y) \mid Z$ | conditional independence |
| $(X \not\perp\!\!\!\perp Y) \mid Z$ | failure of conditional independence |

#### SI Appendix 1: Natural selection requires additive transgenerational effects on fitness at one level.

The Price equation is agnostic about inheritance: it tracks evolutionary change using only fitness $\omega$, phenotype $z$, and the parent–offspring difference $\delta$, without assuming genes, genotypes, or mechanisms of transmission.

$$\Delta\bar{z} = cov(\omega, z) + E[\omega, \delta] \qquad \text{(S1.1)}$$

Nonetheless, rewriting the Price equation in terms of offspring phenotype reveals the same *additive* structure emphasized in Fisher's (1930 ) and Robertson's (1966) theorems. Several authors have developed and demonstrated such reformulations in detail (Frank, 1997, 1998; Queller, 2017). To see the connection requires a minor re-expression of the Price equation in terms of offspring phenotype. Let $z_i^o$ denote the mean phenotype of the offspring of the $i^{th}$ parent as

$$z_i^o = z_i + \delta_i \qquad \text{(S1.2)}$$

Eq. S1.1 can now be used to re-express the mean generational change of offspring phenotype as

$$\Delta\bar{z} = cov(\omega, z^o) + \bar{\delta}_T \qquad \text{(S1.3)}$$

Here, the $\bar{\delta}_T$ term represents the difference in mean phenotypic values across the two generations if there had been no selection (i.e., the non-selective "transmission bias"). The first quantity on the right in Eq. S1.3 is the selective covariance, and it illustrates that what matters to natural selection is the covariance between (*i*) the fitness of the parents and (*ii*) the degree to which fitness effects on parental phenotype can be *transmitted* to the offspring phenotype in the next generation.

The mean phenotype of the offspring of the $i^{th}$ parent can alternatively be written in terms of a regression

$$z_i^o = \bar{z} + \beta_{z^o,z}(z_i - \bar{z}) + \varepsilon_{i_{nil}} \qquad \text{(S1.4)}$$

The residual error $\epsilon_{i_{nil}}$ refers to the assumption that any other causes of phenotypic variation between $z_i^o$ and $z_i$ must be unrelated to fitness of the $i^{th}$ parent.

In Eq. S1.3 $\bar{\delta}_T$ is assumed to be zero, and it is hereafter excluded for simplicity. Given that for any constant (*C*) $cov(x, C) = 0$ and $cov(x, Cy) = C \cdot cov(x, y)$, Eq. S1.3 can be updated as

$$\Delta\bar{z} = \beta_{z^o,z} \cdot cov(\omega, z) \qquad \text{(S1.5)}$$

Since the selective covariance, $cov(\omega, z)$, from Eq. S1.5 is equivalent to the selection differential, *S,* in quantitative genetics (Robertson, 1966), the breeder's equation for the response to selection (*R*) can be derived from the Price equation as follows:

$$\Delta\bar{z} = \beta_{z^o,z} \cdot S \tag{S1.6}$$

$$R = h^2 S \tag{S1.7}$$

The connection between the Price and breeder's equations ($\beta_{z^o,z} = h^2 = V_A/V_P$) highlights how natural selection from *the view of a reproducing organism* requires additive genetic variance ($V_A$) for a trait. Thus, the Price equation recovers the breeder's equation when offspring phenotype is linearly predictable from parental phenotype, with the strength of this mapping captured by $h^2$. However, there can be conditions where additive genetic variance, $V_A$, will not be equivalent to $cov(z^o, z)$, and from the Price equation we understand that it is the latter that really matters for evolution by natural selection.

Using $cov(x, y) = \beta_{y.x} \cdot var(x)$, Eq. S1.5 can be further taken apart.

$$\Delta\bar{z} = \beta_{z^o,z}\, \beta_{\omega,z}\, var(z) \tag{S1.8}$$

The presence of two linear coefficients within this result might be surprising given that the Price equation starts with no mechanistic assumptions about the joint distribution of fitness and phenotype. In other words, because the Price equation does not start with causal mechanisms, it cannot assert that the causal processes are, in fact, linear in nature. What this expression does reveal is that natural selection within a single population can only act on whatever *reliably manifests* in the next generation as an additive effect on the organism's phenotype. Note that this conclusion is specific to organism-level selection, not to multi-level or contextually dependent systems. Interestingly, Fisher's theorem (1930) also can be derived from the Price equation (Frank, 1997), and Fisher emphasized that additive genetic variance was central to understanding the fundamental nature of natural selection. Regardless, a consequence of this equation is that non-additive interactive effects (*e.g.*, IGEs and IIGEs) cannot be the target of selection when natural selection operates only at a single level (*i.e.*, the same conclusion drawn from the heritability criterion of Lewontin's recipe). This underscores why higher-level emergent effects, such as interactive phenotypes or collective mediators, fall outside the scope of classical organism-level selection and require the synaptive framework developed in **SI Appendix 2**.

### SI Appendix 2: Irreducible Cross-Level Covariance and the Limits of MLS1

#### 2.1 Purpose

This appendix establishes the general condition under which the classical MLS1 Price decomposition correctly represents causal group effects and explains why this condition fails in systems where fitness is mediated by emergent, configuration-dependent group traits. **SI Appendix 1** outlined the additive baseline for natural selection; here consequences of relaxing that assumption are examined.

#### 2.2 Causal models and notation

To make this distinction apparent, the Price equation is re-expressed below using causal graphs where fitness is a simultaneous function of individual traits and group-level traits. The concepts and notation follow the causal structure presented in **Figure S2.1**. The figure identifies how an emergent mediator is distinctly different from an additive aggregate.

**Figure S2.1**

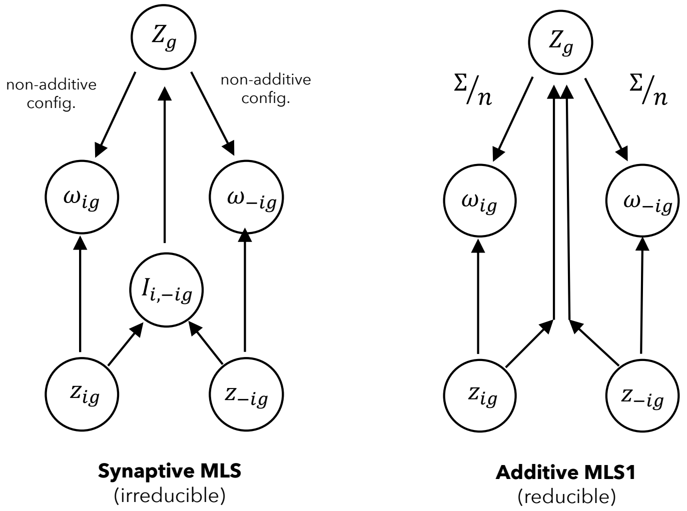

$$Z_g = F(z_{1g}, \ldots z_{ng}) \qquad Z_g = F(z_{1g}, \ldots z_{ng})$$

$$Z_g \not\equiv F(\bar{z}_g) \qquad Z_g \equiv F(\bar{z}_g)$$

In Figure S2.1:

$z_{ig}$ is the trait of individual *i* in group *g*

$z_{-ig}$ is a vector of traits in other individuals in *i*'s group (*g*)

$I_{i,-ig}$ is the interaction that constructs an emergent group trait (*e.g.*, public goods dynamics)

$\omega_{ig}$ is individual fitness as a function of (i) self, (ii) others and (iii) an emergent group trait

$\boldsymbol{Z}_g$ is a group trait:

In synaptive MLS, $\boldsymbol{Z}_g$ is an emergent group trait (*e.g.*, quorum sensing, sanguivory, etc.)

In MLS1, $\boldsymbol{Z}_g$ is a genuine additive cause that "screens off" the lower-level configuration

It is important to distinguish between the variables that mediate fitness causally at the group level (**Fig S2.1**) and the traits used to measure selection in a Price-equation decomposition (Heisler & Damuth, 1987; Damuth & Heisler, 1988). In Price-equation analyses, group-level selection is typically quantified using an additive summary of group composition; denote such a summary by $\theta_g$. The Price equation places no requirement on which traits may be used to measure selection; however, the interpretation of group-level effects depends on how group traits are represented. When a group-level causal mediator $Z_g$ can be summarized additively from individual traits by $\theta_g$, group-level Price covariances will recover its effect. When $Z_g$ instead depends on how individuals are arranged or interact, $\theta_g$ will be insufficient, and Price-based group terms no longer track the causal mechanism shaping fitness. This distinction motivates the irreducibility criterion developed below.

##### 2.3 Synaptive MLS

Rather than being due solely to the individuals own trait, individual fitness under **Synaptive MLS** is expressed as function of a composition-dependent collective environment in which emergent traits arise:

$$\omega_i = \varphi(z_i, z_{-i}, \boldsymbol{Z}_g). \quad \text{(S2.1)}$$

Here, $\varphi$ denotes a causal function that may involve threshold effects, nonlinear gains, or diminishing returns. Thus, *in synaptive systems*, the emergent group trait $\boldsymbol{Z}_g$depends on the configuration of the group: it is not reducible to a sum or mean of individual traits. Substituting this causal structure into the Price equation (Price, 1970) yields a form of selection in which the only selective covariance is the cross-level covariance between individual traits and emergent-mediated fitness:

$$\Delta\bar{z} = cov(\varphi(z_i, z_{-i}, \boldsymbol{Z}_g), z_i) + E[\varphi(z_i, z_{-i}, \boldsymbol{Z}_g), \delta_i]. \quad \text{(S2.2)}$$

#### 2.4 The classical MLS1 decomposition and its implicit assumptions

The classical form of the Price equation for MLS1 (Damuth and Heisler, 1988) describes evolution as a sum of group-level selective covariance (group-specific), individual selective covariance (independent of group), and a transmission bias term:

$$\Delta\bar{z} = cov(\bar{z}_g, \bar{\omega}_g) + E[cov(\omega_i, z_i|g)] + E[\omega_i, \delta_i]. \quad \text{(S2.3)}$$

where $\bar{z}_g = E[z_i \mid g]$ is an additive group predictor. This is a causally appropriate decomposition only if the group mediator $Z_g$ is *identical in information content* to $\bar{z}_g$:

$$Z_g \equiv F(\bar{z}_g). \quad \text{(S2.4)}$$

This is the **MLS1 additivity condition**. If an emergent causal trait $Z_g$ cannot be written as a function of $\bar{z}_g$, then the MLS1 group term cannot represent the causal group effect (Fig S2.1).

#### 2.5 Conditional independence and reducibility to MLS1 levels of selection

The most common additive summary in the classical MLS1 decomposition is the group mean $\bar{z}_g$. Depending on the system, other relevant summaries include: a sum, a frequency, a weighted average, or some linear combination of summaries. Hence, $\theta_g$ is used rather than $\bar{z}_g$ to emphasize that MLS1 requires only that *some* additive group to eliminates the dependence of the emergent group mediator on individual traits.

Under the MLS1 additivity condition, the detailed individual composition of the group does not matter to the between-group covariance in Eq. S2.3. This condition is equivalent to the statement that $Z_g$ is conditionally independent of individual traits $(z_i, \ldots, z_{ng})$ once the additive predictor $\theta_g$ is known:

$$(Z_g \perp\!\!\!\perp z_i) \mid \theta_g \quad \text{(S2.5)}$$

where $\theta_g = \bar{z}_g$ or any other additive summary.

This means that that once a group trait is fully determined by some additive summary of the individuals (*e.g.*, group mean trait $\bar{z}_g$), then

(i) the emergent trait $Z_g$ adds no further information about fitness beyond that additive summary, and

(ii) individual fitness does not depend on the group configuration of the trait values of individual in the group.

2.5 guarantees *reducibility* to the decomposition given by Eq. S2.3. If Eq. S2.5 holds then for all individuals in group *g*:

$$\varphi(z_i, z_{-i}, Z_g) = \varphi(z_i, \theta_g). \tag{S2.6}$$

Thus, under conditional independence, the emergent group mediator $Z_g$ becomes functionally equivalent to its additive summary. Therefore:

$$cov(Z_g, \bar{\omega}_g) = cov(\theta_g, \bar{\omega}_g). \tag{S2.7}$$

**2.6 Irreducible cross-level dependence: MLS1 vs. Synaptive MLS**

The additivity assumption required by MLS1 will be violated when $Z_g$ is a non-additive function of the trait distribution (*e.g.*, depends on variance, coexistence, thresholds, joint presence of types). Thus, MLS1 will fail when:

$$Z_g \not\equiv F(\theta_g), \qquad (Z_g \not\perp\!\!\!\perp z_i) \mid \theta_g. \tag{S2.8}$$

In this case,

$$\varphi(z_i, z_{-\mathrm{i}}, Z_g) \neq \varphi(z_i, \theta_g), \tag{S2.9}$$

and reducing covariance to the MLS1 levels of selection (Eq. S2.3) is a distortion of the true group-level causal influence.

If Eq. S2.8 holds, the emergent group trait retains information about fitness that cannot be "screened off" by any additive group predictor $\theta_g$. This represents *irreducible cross-level covariance*, the defining signature of synaptation.

Recall that the *synaptive fitness mapping* was defined as: $\omega_i = \varphi(z_i, z_{-\mathrm{i}}, Z_g)$

Let the *additively reduced fitness mapping* be defined as:

$$\omega_i^{\theta} = \varphi(z_i, \theta_g). \quad (S2.10)$$

Formally, synaptive covariance is present when:

$$cov(\omega_i, z_i) \neq cov(\omega_i^{\theta}, z_i), \quad (S2.11)$$

for all additive $\theta_g$.

#### 2.7 Summarizing three kinds of group effects

Section 2.3 described the group effect that is due to irreducible selective covariance. Section 2.4 described a second kind of group effect, that is due to a fully additive predictor that "screens off" the lower-level configuration of the group. Section 2.6 described how the MLS1 decomposition misrepresents the genuine causal group effect of the first kind (irreducible selective covariance). Okasha (2006) identified a third effect, which manifests as a false MLS1 group covariance when no group-level causal variable exists. Okasha (2006) refers to this artifact as a *cross-level by-product* of selection on lower-level individual organisms. Once the individual traits are conditioned upon, the artifactual group-level effect disappears. Table below summarizes these cases.

| Category | Causal status | Reducibility conditions | Example |
|---|---|---|---|
| **MLS1 group effect** | Additive; "screens off" lower-level configuration | *reducible* $Z_g \equiv F(\bar{z}_g)$ $(Z_g \perp\!\!\!\perp z_i) \mid \bar{z}_g$ | Mean toxin-degrading enzyme concentration predicts how ambient toxin levels affect fitness. |
| **Cross-level by-product** | No group-level cause: group effect is artifact | *Trivially reducible* no $Z_g$ exists; conditioning on individual traits eliminates group-level covariance. | Groups with more high-fertility individuals produce more offspring; (*e.g.*, Figure 2 in main text) |
| **Synaptive group effect** | Emergent; cross-level entanglement. | *irreducible* $Z_g \not\equiv F(\bar{z}_g)$ $(Z_g \not\!\perp\!\!\!\perp z_i) \mid \bar{z}_g$ | Division of labour: complementary types must co-occur for high fitness. |

#### SI Appendix 3. A Minimal Numerical Example of Irreducible Synaptive Covariance

This appendix presents a compact numerical example showing how synaptive MLS1 generates a selective covariance term that cannot be reduced to independent within- and between-group effects. The example is intentionally minimal to highlight the causal structure identified in **SI Appendix 2**.

##### 3.1 Setup

Individuals belong to one of two groups, $g$. Each individual carries a scalar trait value $z_i$. Define two group statistics mediated by an emergent group trait:

$$Z_g = \frac{1}{n_g} \sum_{i \in g} z_i,$$

$$Z_g^{(2)} = \frac{1}{n_g} \sum_{i \in g} z_i^2 .$$

Let individual fitness be determined by **cross-level interaction** between an individual effect ($z_i^2$) and a group effect ($Z_g$):

$$w_i = z_i^2 Z_g.$$

Then, in this simple causal model, all group-level variation in $\bar{w}_g$ arises from the term $Z_g^{(2)}\, Z_g$:

$$E[w_i \mid g] \;=\; Z_g^{(2)}\, Z_g.$$

##### 3.2 Model: Two task groups with identical means but different compositions

Consider two groups, each with three individuals:

| Task Group | Trait values ($z_i$) | Mean ($Z_g$) | Second moment ($Z_g^{(2)}$) |
|---|---|---|---|
| A | 0, 1, 3 | 2 | 5 |
| B | 1, 1, 2 | 2 | 3 |

Both task groups have the same mean trait:

$$Z_A = Z_B = 2,$$

but they differ in their second crude moment:

$$Z_A^{(2)} = 5, \;\; Z_B^{(2)} = 3.$$

The group mean fitness values are

$$\bar{w}_A = Z_A^{(2)} Z_A = 10,$$
$$\bar{w}_B = Z_B^{(2)} Z_B = \; 6.$$

Note that even though the task groups A and B have identical means, their expected fitness differs by ~67% (as an increase from 6) because their cross-level quantities differ. This difference will be "invisible" to any model that treats $Z_g$ as a causal variable that "screens off" the composition of the group (**SI Appendix 2**).

**3.3 Between-group covariance**

The between-group selective covariance in the classical form of the Price equation for MLS1 is:

$$cov_g(E[w_i \mid g], E[z_i \mid g]) = cov_g\left(Z_g^{(2)} Z_g \,, Z_g\right).$$

Because $Z_g$ does not vary across groups ($Z_g \; = 2$), the covariance $cov_g\left(Z_g \,, \overline{w}_g\right)$ is necessarily zero.

$$cov_g\left(Z_g \,, \overline{w}_g\right) = \frac{1}{2}(0 + 0) = 0.$$

Yet there exists a real group-level effect on mean fitness, driven by the difference in $Z_g^{(2)}$.

$$cov_g\left(Z_g^{(2)} \,, \overline{w}_g\right) = \frac{1}{2}(2 + 2) = 2 \; > 0.$$

So, this minimal numerical example shows:

- The **Price "group term", when written with a** $Z_g$ as a causal variable will "screen off" the composition of the group and report *no* between-group covariance.

- The **true** group-level component of the fitness function is $Z_g^{(2)} Z_g$, with all the variation in $\bar{w}_g$ is coming from $Z_g^{(2)}$, not $Z_g$.

Thus, for any function ($f$) of the group mean alone, the selective covariance is not reducible to individual (MLS1 fitness) or group (MLS2 fitness) decompositions, or any additive combinations of the two:

$$cov_g\left(Z_g^{(2)} Z_g , Z_g\right) \neq f\left(Z_g\right).$$

Group means, alone, cannot represent how internal group organization modulates fitness.

##### 3.4 Summary

This covariance is **irreducible**: it depends jointly on $Z_g$ and $Z_g^{(2)}$. Two groups share the same mean $Z_g$ while differing in $Z_g^{(2)}$, so no function $f\left(Z_g\right)$ can reproduce $\bar{w}_g$. Even in this minimal system, cross-level causal interaction produces fitness differences that cannot be detected by any covariance involving a single additive group descriptor. This is the signature of synaptive covariance.

##### 3.5 Implication 1: Failure of simple covariance decompositions

This example exposes a general limitation of covariance decompositions in systems where fitness depends on configuration-dependent interactions. Although there is a genuine causal group effect here (differences in $Z_g^{(2)}$ produce large differences in $\bar{w}_g$), both the classical covariance $\mathrm{Cov}_g(\bar{z}_g, \bar{w}_g)$ and the covariance involving $Z_g$ failed to detect it. The problem is structural: when fitness depends jointly on multiple emergent quantities, no single scalar group summary $\theta_g$ (mean, sum, frequency, or any linear transformation) will yield a covariance $\mathrm{Cov}_g(\theta_g, \bar{w}_g)$ that recovers the magnitude or even the sign of the group-level effect. Because the group effect arises from the interaction of emergent quantities, any one-dimensional covariance projection could cancel or mask the underlying causal influence. Thus, there is no simple covariance decomposition that reliably indicates the presence or direction of the group-level effect of selection.

##### 3.6 Implication 2: How to diagnose synaptation

This covariance failure motivates the diagnostic method developed in **SI Appendix 5**. This example illustrates that the appropriate question is not "*which covariance captures group selection?*" but rather "*can the effect of the emergent mediator be screened off by any additive group descriptor?*" A diagnostic should evaluate this by testing whether any additive predictor $\theta_g$ eliminates the contribution of an emergent mediator $E_g$, and by assessing the conditional-independence criterion $\left(E_g \perp\!\!\!\perp z_i\right) \mid \theta_g$. Failure of screening-off indicates an irreducible synaptive group effect. Thus, **SI Appendix 3** illustrates how simple covariance decompositions can be misleading, and highlights the need for a conditional-independence diagnostic for detecting synaptation.

#### SI Appendix 4. A Biologically Intuitive Example: Division-of-Labour Synaptation

The minimal example presented in **SI Appendix 3** algebraically demonstrates irreducibility. The following scenario shows how the same causal structure can arise in a simple biological context. In this scenario a collective task is carried out by a group of independently replicating entities that perform different roles, a *task group*. This example illustrates how group-level selective covariance can arise through an emergent division-of-labour mediator, $Z_g$, even though groups have no group-level mechanism of heredity.

##### 4.1 Roles and emergent mediator

Two roles are performed by lower-level entities having independent hereditary mechanisms:

- Role 1 (R1), with $z_i = 0$
- Role 2 (R2), with $z_i = 1$

Define a non-additive emergent group mediator:

$$Z_g = \begin{cases} 1 & \text{if group } g \text{ contains at least one R1 and at least one R2,} \\ 0 & \text{otherwise.} \end{cases}$$

Thus, $Z_g$ represents a minimal "division of labour" property. It cannot be written as any additive summary of the individual $z_i$.

##### 4.2 Fitness

When $Z_g = 0$, all individuals have fitness 1.

When $Z_g = 1$, all individuals benefit but unequally:

$$w_i = \begin{cases} 1.5 & \text{if } Z_g = 1 \text{ and lower-level entity } i \text{ carries out R1,} \\ 1.2 & \text{if } Z_g = 1 \text{ and lower-level entity } i \text{ carries out R2,} \\ 1 & \text{if } Z_g = 0. \end{cases}$$

When a group attains division-of-labour ($Z_g = 1$), all group members benefit, but those carrying out Role 1 benefit more than those carrying out Role 2. The asymmetric benefit of the division of labour is used to illustrate the need for a synaptive covariance term.

##### 4.3 Group composition and fitness

Consider four groups, each with just three members, with composition and fitness as follows.

| **Group** | **Composition** | **Mediator** ($Z_g$) | **Individual fitness** ($w_i$) | **Group mean fitness** ($\bar{w}_g = E[w_i \mid g]$) |
|---|---|---|---|---|
| **G1** | R1=3, R2=0 | 0 | 1, 1, 1 | $\bar{w}_1 = 1.0$ |
| **G2** | R1=2, R2=1 | 1 | 1.5, 1.5, 1.2 | $\bar{w}_2 = (1.5 + 1.5 + 1.2)/3 = 1.4$ |
| **G3** | R1=1, R2=2 | 1 | 1.5, 1.2, 1.2 | $\bar{w}_3 = (1.5 + 1.2 + 1.2)/3 = 1.3$ |
| **G4** | R1=0, R2=3 | 0 | 1, 1, 1 | $\bar{w}_4 = 1.0$ |

Means across groups: $\bar{Z} = 0.5,\ \bar{w} = 1.175.$

Note how the division-of-labour state **(**$Z_g$**)** alters the distribution of fitness *within* each group ($\bar{w}_2 = 1.4, \bar{w}_3 = 1.3, \bar{w}_{1,4} = 1.0.$). The emergent mediator creates systematic differences in group mean fitness. Because groups vary in $Z_g$, the between-group covariance will be nonzero. Importantly, in this example, the effect of the mediator $Z_g$ cannot be determined from the individual traits: no weighting of the individual $z_i$ values will return the mediated group fitness $\bar{w}_g$ by aggregation. Furthermore, it's possible to construct groups with the same mean trait, $\bar{z}_g$, that differ in $Z_g$. Thus, this scenario fails to meet conditional independence, which is required by the classical form of the Price equation for MLS1 (**SI Appendix 2**). In this scenario:

$$Z_g \not\perp\!\!\!\perp z_i \mid \bar{z}_g.$$

##### 4.4 Synaptive covariance verses MLS1 covariance

Because the emergent group property $Z_g$ contains relational information unavailable from $\bar{z}_g$, the differences in group fitness are irreducible. Given that the emergent mediator creates systematic differences in group mean fitness, the covariance with the emergent mediator is positive:

$$cov_g(Z_g, \bar{w}_g) = \frac{1}{4}\sum_{g=1}^{4}(Z_g - \bar{Z})(\bar{w}_g - \bar{w}) = 0.0875 > 0.$$

In contrast, a covariance can be computed according to the additive MLS1 group trait $\bar{z}_g$. This approach treats selection as if it were acting on an additive function of the role value; effectively imposing a fictitious mapping $w_i = \varphi(z_i, \bar{z}_G)$. This covariance, which cannot capture relational structures such as division-of-labour, dilutes and reverses the sign of the group-level selective covariance:

$$cov_g(\bar{z}_G, \bar{w}_G) = \frac{1}{4}\sum_{G=1}^{4}(\bar{z}_G - \bar{z})(\bar{w}_G - \bar{w}) \approx -0.00417$$

Although $\bar{w}_g$ can be rewritten algebraically as a piecewise function of the group mean $\bar{z}_g$, this does not identify $\bar{z}_g$ as the causal group-level determinant of fitness. The relevant causal mediator is the emergent trait $Z_g$, which encodes the configuration-dependent structure associated with division of labour and cannot be recovered from any additive summary of individual traits. The covariance with $\bar{z}_g$ consequently misrepresents both the magnitude and direction of the group effect. This demonstrates that the additive MLS1 decomposition is not aligned with the underlying causal architecture, and motivates the need for a diagnostic test to assess whether an additive group descriptor can screen off the emergent mediator or whether the system instead manifests irreducible synaptive covariance.

##### 4.5 Conclusion

In this example, the task group has no group-level mechanism for heritability**.** Yet there exists irreducible selective covariance at the group level. The selected effect is enabled by a division of labour that cannot be captured by any additive summary of the individual traits, even though fitness can be algebraically rewritten as a function of the mean. This identifies a clear case of a **synaptation**: a selected effect enabled by cross-level causal structure rather than group reproduction. Applying the classical MLS1 form of the Price equation in this case masks or reverses the group effect because it is not aligned with the casual architecture of fitness.

**SI Appendix 5: A Diagnostic for Synaptation (Irreducible Cross-Level Selective Covariance)**

This appendix presents a proposal for an empirical method for diagnosing synaptation, which is defined as cases in which individual fitness depends on an emergent group-level mediator whose causal influence cannot be screened off by any additive summary of group composition. The diagnostic operationalizes the irreducibility criterion developed in **SI Appendix 2**. It builds directly on the regression-based contextual analysis framework developed by Heisler and Damuth (1987), extending it beyond additive partitions by testing for the failure of conditional independence between individual traits and an emergent group-level mediator after conditioning on all additive group descriptors.

Note that this diagnostic does not assume that a Price-equation group covariance will recover the causal group effect; rather, it directly tests the condition under which such recovery would be possible (i.e., whether any additive summary can screen off the emergent mediator). The diagnostic is hypothesis-guided rather than exploratory: it is designed to test whether a theoretically motivated candidate for an emergent mediator exhibits irreducible cross-level mediation, not to discover emergent mediators through data exploration alone.

The diagnostic distinguishes three conceptually distinct cases:

1. Cross-level by-products (false additive group effects, *sensu* Okasha 2006),
2. Genuine but additively representable group effects, and
3. Irreducible synaptations, characterized by configuration-dependent causal entanglement.

When multiple plausible candidates for $Z_g$ exist, the diagnostic can be applied sequentially or in parallel with appropriate correction, but failure of additive screening for any candidate is sufficient to establish irreducible cross-level selective covariance in the system.

##### 5.1 Data structure and notation

Individuals $i = 1, \dots, N$ are nested within groups $g = 1, \dots, G$.

- $z_i$: focal individual trait
- $w_i$: individual fitness
- $A_g$: A vector of additive group descriptors, including any summary of group composition that is additive in individual traits (e.g. means, sums, frequencies, additive moments, or linear combinations thereof).

- $C_g$: A vector of configuration-dependent descriptors, capturing aspects of group structure that depend on arrangement, co-occurrence, or interaction topology rather than additive totals (e.g., role heterogeneity, complementarity indices, network structure).
- $Z_g$: An emergent group mediator, a collective property hypothesized to causally influence fitness and may or may not be representable by an additive summary. $Z_g$ is the target of the screening-off test.

Crucially, $Z_g$ is never assumed to be additive. It may be additively representable in some systems, but this is an empirical question addressed by the diagnostic.

##### 5.2 STEP 1: Baseline contextual analysis

Start with the standard contextual analysis (Heisler & Damuth, 1987)) used in MLS1:

$$w_i = \alpha + \beta_{\text{ind}} z_i + \beta_{\text{grp}} \bar{z}_g + \varepsilon_i \qquad \text{S5.1}$$

where $\bar{z}_g$ is the group mean of the focal trait.

A nonzero $\beta_{\text{grp}}$indicates between-group selective covariance but does not establish a genuine group-level cause. Such false signal may arise as a cross-level by-product of additive composition (Okasha, 2006; Fig 1 in main text).

If no between-group effect is detected, the diagnostic terminates. Termination here reflects the absence of detectable group-level selection under the chosen scaling, not the absence of configuration dependence or emergent structure per se.

##### 5.3 STEP 2: Additive screening using $A_g$

Test if the between-group effect is reducible to additive composition, by fitting:

$$w_i = \alpha + \beta_{\text{ind}} z_i + \gamma^{\top} A_G + \varepsilon_i \qquad \text{S5.2}$$

The vector $A_g$ should include all plausible additive summaries of group composition relevant to the system. In practice, $A_g$ should include means, sums, frequencies, and other additive moments of individual traits that plausibly contribute to the emergent mediator $Z_g$. The specification of $A_g$ should be justified through domain knowledge.

The term $\gamma^{\top} A_g$ represents the combined influence of all additive features of group composition on an individual's fitness, where $\gamma^{\top}$ specifies how strongly each additive feature contributes. In other words, $\gamma^{\top} A_g$ describes what can be predicted from who is in the group, without considering how group members interact or organize.

Diagnostic question:

Does there exist some specification of $A_g$ such that the apparent group-level effect is fully accounted for?

- If yes, the effect is an additive cross-level by-product in Okasha's sense.
- If no, additive reducibility fails and the analysis proceeds.

##### 5.4 STEP 3: Configuration-dependent predictors $\boldsymbol{C_g}$

Next, assess whether configuration-dependent structure explains the remaining group effect by fitting:

$$w_i = \alpha + \beta_{\text{ind}} z_i + \gamma^{\top} A_g + \eta^{\top} C_g + \varepsilon_i \qquad \text{S5.3}$$

This step determines whether the group effect depends on interaction structure rather than mere aggregation. It operationalizes the algebraic failures of additive screening illustrated in SI Appendix 3 and the biological division-of-labour example in SI Appendix 4 by testing whether configuration-dependent structure explains fitness variation beyond additive composition.

The term $\eta^{\top} C_g$ represents how group organization influences individual fitness, where $\eta^{\top}$ specifies how strongly different aspects of group configuration (*e.g.*, role differentiation, interaction patterns, or coordination) affect fitness.

The purpose of Step 3 is to test whether configuration matters at all to fitness. Step 3 does not, by itself, diagnose synaptation. Synaptation is diagnosed only when fitness depends on configuration *and* that dependence cannot be reduced to an additive descriptor of group composition. Step 4 introduces $Z_g$ as an explicit candidate emergent mediator on the causal pathway from configuration to fitness, allowing direct testing of whether the configuration remains causally relevant and fails additive screening.

A nonzero $\eta^{\top} C_g$ therefore establishes configuration dependence, but does not imply irreducibility. This corresponds to the failures in SI Appendix 3 and the division-of-labour mechanism in SI Appendix 4, where relational structure, not additive totals, determines fitness.

#### 5.5 STEP 4: Emergent mediator $Z_g$ and conditional independence

The tests used below are intended to detect statistical dependencies that are diagnostic of the proposed causal architecture; full causal identification would require further assumptions or experimental intervention.

Next the emergent group mediator is introduced:

$$w_i = \alpha + \beta_{\text{ind}} z_i + \gamma^\top A_g + \delta Z_g + \varepsilon_i. \qquad \text{S5.4}$$

Two tests are required.

**STEP 4.1: Screening of the causal effect on fitness**

Test if the mediator retains explanatory power after conditioning on all additive summaries:

$$H_0: \delta = 0 \text{ vs. } H_A: \delta \neq 0.$$

If $\delta$ cannot be eliminated by specifying an additive summary $A_g$, then the causal influence of $Z_g$ cannot be screened off.

**STEP 4.2: Conditional independence test (core diagnostic)**

Under additive representability, the emergent mediator must satisfy:

$$Z_g \perp\!\!\!\perp z_i \mid A_g. \qquad \text{S5.5}$$

Eq. S5.5 should be interpreted as a sufficiency condition on additive group descriptors. It states that once all additive information about group composition is captured by $A_g$, the emergent mediator $Z_g$ should have no additional dependence on individual traits. Failure to meet this condition indicates that $Z_g$ encodes configuration-dependent information that cannot be reduced to any additive summary of individual traits. This expresses the MLS1 additivity condition.

Eq. S5.5 can be tested using partial correlations. Such tests assess whether the correlation between $Z_g$ and $z_i$ remains nonzero after statistically controlling for all components of

$A_g$. Under additive representability, conditioning on $A_g$ should eliminate any association between individual traits and the emergent mediator; a residual partial correlation therefore indicates failure of additive screening.

Partial-correlation tests assume linear relationships. Other testing frameworks such as conditional mutual information or non-parametric graph-based methods may be more appropriate in systems where nonlinear dependencies are expected. While the precise statistical implementation may vary, the criterion is sufficient to diagnose irreducible cross-level mediation.

Failure of Eq. S5.5 indicates that no additive group descriptor screens off the emergent mediator. This is the defining empirical signature of irreducible synaptive covariance.

##### 5.6 STEP 5: Composition randomization

To confirm that the detected effect depends on configuration rather than aggregation, group membership is randomized while preserving the marginal distribution of individual traits.

The purpose of the randomization step is to test whether fitness-relevant group effects depend on naturally assembled configuration rather than aggregation alone. Randomization may disrupt the effect either by breaking the relationship between $Z_g$ and fitness or by preventing the construction of $Z_g$ itself; in both cases, loss of the signal indicates configuration-dependent mediation.

The diagnostic steps (Eqs. S5.1–S5.5) are recomputed on randomized data. Following permutation, $Z_g$is recalculated for the permuted groups before re-running Eqs. S5.1–S5.5. The specific randomization scheme should preserve structural features relevant to the hypothesis under test (e.g., group size or role structure), and its precise implementation is necessarily context-dependent and deferred to future applications.

A synaptive signal must:

- disappear under randomization, and
- reappear in naturally assembled groups.

This step guards against spurious inference due to sampling structure.

##### 5.7 Decision rules

The collective trait mediated by $Z_g$ is classified as a synaptation if and only if all the following hold:

1. Between-group selective covariance exists (Eq. S5.1).
2. No additive summary $A_g$ eliminates the group effect (Eq. S5.2).
3. Configuration-dependent predictors do not render the effect additively reducible (Eq. S5.3).
4. The emergent mediator $Z_g$ retains causal influence on fitness after additive controls (Eq. S5.4).
5. Conditional independence fails:

$$(Z_g \not\perp\!\!\!\perp z_i) \mid A_g \quad \text{S5.6}$$

6. Randomization destroys the signal and natural configuration restores it.

Together, these establish irreducible cross-level selective covariance, a hallmark of synaptation.

##### 5.8 Relation to causal diagrams

In causal-graph terms, additive reducibility corresponds to $A_g$ blocking all paths from individual traits to the emergent mediator and from the mediator to fitness. Synaptation is diagnosed when no additive node blocks these paths, leaving cross-level causal pathways unshielded (**SI Appendix 2**).
